# Bump attractor dynamics underlying stimulus integration in perceptual estimation tasks

**DOI:** 10.1101/2021.03.15.434192

**Authors:** Jose M. Esnaola-Acebes, Alex Roxin, Klaus Wimmer

**Affiliations:** Centre de Recerca Matemàtica, Campus de Bellaterra, Edifici C, 08193 Bellaterra (Barcelona), Spain; Barcelona Graduate School of Mathematics, Barcelona, Spain

## Abstract

Perceptual decision and continuous stimulus estimation tasks involve making judgments based on accumulated sensory evidence. Network models of evidence integration usually rely on competition between neural populations each encoding a discrete categorical choice and do not maintain information that is necessary for a continuous perceptual judgement. Here, we show that a continuous attractor network can integrate a circular stimulus feature and track the stimulus average in the phase of its activity bump. We show analytically that the network can compute the running average of the stimulus almost optimally, and that the nonlinear internal dynamics affect the temporal weighting of sensory evidence. Whether the network shows early (primacy), uniform or late (recency) weighting depends on the relative strength of the stimuli compared to the bump’s amplitude and initial state. The global excitatory drive, a single model parameter, modulates the specific relation between internal dynamics and sensory inputs. We show that this can account for the heterogeneity of temporal weighting profiles and reaction times observed in humans integrating a stream of oriented stimulus frames. Our findings point to continuous attractor dynamics as a plausible mechanism underlying stimulus integration in perceptual estimation tasks.

---

Integrating information over time is a fundamental computation that neural systems need to perform in perceptual decision and continuous estimation tasks. While a categorical perceptual decision usually requires discriminating stimuli in order to select between several options (e.g. left vs. right motion of an ambiguous random dot stimulus^1^, or clockwise vs. counterclockwise motion compared to a reference^2^), in stimulus estimation tasks participants report a continuous stimulus feature, for example the stimulus orientation in degrees. Typical estimation tasks involve accumulating evidence about the average motion direction of a random dot pattern in degrees^2–5^ or the average of a sequence of oriented gratings^6,7^. In particular, we are interested here in experimental tasks that require estimating, as an analog quantity, the temporal average of a circular feature such as the orientation or direction of a potentially time-varying stimulus.

Neural network models of evidence integration usually rely on slow recurrent dynamics and competition between neural populations, each encoding one of the categorical choice options^8–11^. The architecture of these *discrete* attractor models thus directly reflects the categorical nature of the decision task. Similarly, phenomenological drift-diffusion models linearly integrate stimulus evidence for or against different choice options until one of the bounds is reached^12–17^. While the amount of accumulated evidence (i.e., the state of the decision variable) at the time of the decision is predictive of choice certainty or confidence^18^, by design these models do not maintain information about continuous features of the integrated stimulus (e.g. the average orientation; but see ref.^19^). They are thus not well suited to study stimulus estimation tasks that require continuous perceptual judgements or tasks that require both the report of a continuous estimate and a categorical choice^2,20,21^.

The representation of continuous stimuli in neural population codes has been studied extensively using normative models^20^ or neural coding models^2,22^. These models can explain how to optimally represent a stimulus in a neural population (e.g. an orientated stimulus in area V1), how to compute an estimate from a noisy sensory input^23^, how to combine different cues in a single representation^24^, and how to read-out the population code in order to optimally discriminate stimuli^25^. These theoretical studies have focused on investigating the representation of stimulus features in sensory cortices rather than the computations involved in perceptual decision making and stimulus estimation. There is a lack of biophysical neural network models that can be used to study the underlying circuit mechanisms of these computations.

A natural candidate for such aneural network model is the well-known *continuous* attractor model^26,27^. Continuous bump attractor networks have a topological ring architecture and due to strong recurrent coupling they can show localized activity in a subset of neurons with a bell-shaped pattern (bump) that persists even in the absence of tuned external input. Because all states along the ring are energetically equal, the bump can be located at any location along the network. The bump location thus provides a substrate for encoding a continuous variable such as an orientation or a spatial location. Moreover, while intrinsically the bump position is stable, external inputs can drive the bump to move. Due to these features, bump attractor networks have been proposed as the neural mechanism underlying a variety of sensory and cognitive computations: the emergence of contrast-invariant orientation tuning in visual cortex ^28^, the impact of microstimulation on the identification of the direction of a weak motion stimulus ^29^, the maintenance of spatial working memory in prefrontal cortex^30,31^, and the representation of spatial orientation and angular path integration in the mammalian head direction system^32–35^, the fly’s heading direction system^36^ and in grid cells^37^. None of these previous studies has investigated whether a bump attractor network can accumulate sensory information to estimate the time-average of a continuous stimulus as required in perceptual estimation tasks.

Here, we show that the ring attractor model can accurately compute the average of a time-varying circular stimulus feature. Close to the dynamical regime where the bump of activity emerges the network approximates a *perfect vector integrator* such that the phase of the bump of activity at each time point corresponds to the running circular average of the stimulus. The temporal weighting of stimulus information is uniform in this case, as is characteristic of perfect integration. Moreover, the model can also exhibit primacy and recency regimes, i.e., it can integrate the stimulus giving higher weight to early or late parts of the stimulus in a fixed duration experiment. These integration regimes depend on the global excitatory drive to the network which is a single parameter controlling the amplitude and the dynamics of the bump. On average, the model estimates the mean stimulus orientation faithfully albeit with a higher trial-to-trial variability when the temporal weighting is not uniform. To illustrate the relevance of our theoretical results, we analyze data from psychophysical experiments in which human observers integrated a stream of eight oriented stimulus frames. We show that the continuous attractor model can quantitatively fit the observed heterogeneity of temporal weighting regimes across subjects and their relationship to the subjects’ reaction times. Our work suggests continuous attractor dynamics as a potential mechanism underlying stimulus integration in perceptual estimation tasks.

## Results

### Stimulus integration with bump attractor dynamics

We start by simulating the integration of stimuli in a continuous ring attractor model, as in experimental tasks that require estimating, as an analog quantity, the average direction of motion of a noisy random dot stimulus^2,3^ or of a sequence of oriented Gabor patches^6,7^. The network model has a ring architecture and Mexican-hat connectivity (Fig. 1a) and its firing rate dynamics are described by Eq. 4 (Methods). Due to strong recurrent connectivity, a bump of activity emerges in the network at a position determined by the stimulus, and this bump persists when the stimulus is removed (Fig. 1b). In this regime, a fast change of the stimulus to a new direction does not, in general, lead to an instantaneous extinction and re-appearance of the bump at the new direction, but rather initiates a continuous translation of the activity along the ring network, directed towards the new stimulus direction (Fig. 1b, Extended Data Fig. 1). This has been termed “virtual rotation”^28^. Thus, the bump position at a given time depends on the current input to the network and on the previous bump position.

**Fig. 1.**
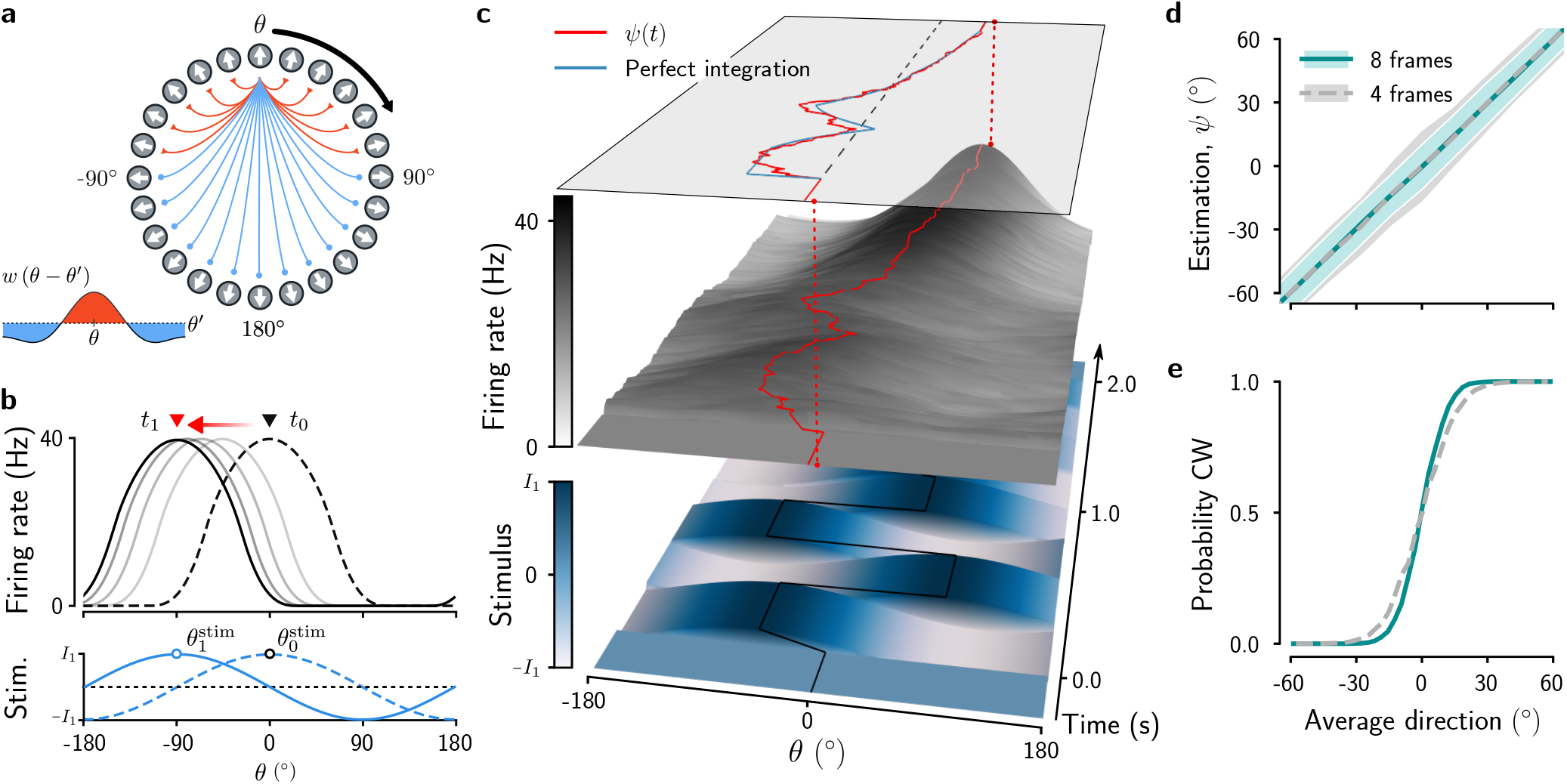
Stimulus integration in the bump attractor network. **a**, Ring network with Mexican-hat connectivity, that is, strong local excitation (red connections) and broader inhibitory coupling (blue). **b**, Network activity for neurons arranged according to their position in the ring (**a**). Due to strong recurrent connectivity, a bump of activity emerges in this network at a position determined by the external input and persists when the input is removed. For time-varying inputs, the activity bump (dashed line) moves towards a new position (solid black line). **c**, Network activity in a single trial. The initial activity of the network (middle panel) is spatially homogeneous and evolves into a bump while integrating a time-varying stimulus composed of eight oriented stimulus frames (bottom panel). Top: The phase of the bump (red) as a function of time closely tracks the running average of the orientations of the stimulus frames (blue). **d**, **e**, Continuous stimulus estimate (**d**) and probability of clockwise choices (**e**) as a function of the average stimulus direction for stimulus durations of 1 s (4 stimulus frames of 250 ms) and 2 s (8 stimulus frames). Categorical choices were obtained by converting positive angles to clockwise reports and negative angles to counterclockwise reports.

We wondered whether through this mechanism the position (phase, *ψ*) of the activity bump could track the average of a time-varying stimulus. We simulated a task which required estimating the average direction of eight successive oriented stimulus frames with constant strength (e.g. stimulus contrast), and directions distributed between −90° and +90° (Fig. 1c). We found that the transient population response to changing stimulus input effectively computes this estimation (Fig. 1c). In the example trial, the evolution of the bump phase closely approximated the cumulative running average of the stimulus, that is, the time-average of the stimulus up to a given time point (Fig. 1c, top). Indeed, estimation curves, obtained by simulating many trials, show that, on average, the bump phase closely tracks a continuous estimate of the averaged sensory input (Fig. 1d). The estimation accuracy improves with the stimulus duration, as expected from an integration process (errorbars in Fig. 1d). This improvement can also be seen in psychometric curves obtained by converting the analog estimate to a categorical choice (Fig. 1e). In sum, these simulations show that the bump attractor network can integrate the stimulus over times much larger than the intrinsic time constant of the network (*τ* = 20 ms, stimulus duration *T*_stim_ = 2 s).

### Dynamics of the bump

Motivated by the observation that the phase of the bump can track the average of the stimulus with striking precision (Fig. 1), we next sought to identify and characterize the neural network mechanisms underlying stimulus integration in the bump attractor network. To study the dynamics of the movement of the bump, we applied a standard perturbation method (Methods) and reduced the network model (Eq. 4) to a 2-dimensional differential equation for the amplitude, *R*(*t*), and the phase of the bump, *ψ*(*t*):

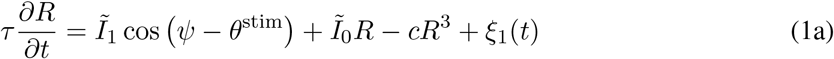

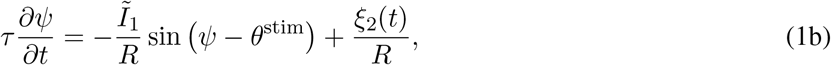

where *τ* is the neural time constant and 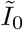 is proportional to the global excitatory drive to each neuron, relative to its critical value at the onset of the bump (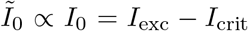; Methods). The constant *c* depends on the synaptic connectivity profile and the nonlinear neural transfer function (Eq. 5). The time-varying stimulus input is characterized by its strength 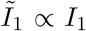, and its orientation *θ*^stim^(*t*). The noise terms *ξ*_1_(*t*) and *ξ*_2_(*t*) are related to the internal stochasticity and to fluctuations in the stimulus realized as independent noise inputs to each neuron in the full model (Eq. 4). The reduced model allows us to study the dynamics of the bump analytically. From Eq. 1b it directly follows that the rate of change of the phase (the angular speed *dψ*/*dt*) depends on the ratio of the stimulus strength *I*_1_ and the bump amplitude *R*. Thus, for a given stimulus strength *I*_1_, the higher the amplitude of the bump the smaller will be the angular speed, that is, larger bumps are more sluggish (Extended Data Fig. 1). The steady-state bump amplitude is determined by the global excitatory drive *I*_0_, (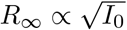; Methods), indicating the key role of this parameter for the integration dynamics.

A deeper understanding about how a time-varying stimulus impacts the phase of the bump can be gained if we visualize the dynamics of the system, Eq. 1, by means of the physical idea of the potential energy. We can interpret the dynamics of the bump as a heavily damped particle sliding down the walls of a 3-dimensional potential well, where a potential Θ (*R, ψ*) of the system described in Eq. 1 is given by

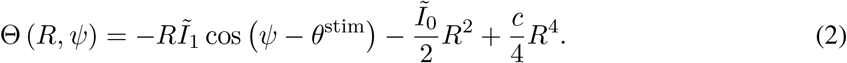

The surface of this potential represents the dynamical manifold of Eq. 1, such that a point on this surface is a possible state of the activity bump, characterized by its amplitude *R* and phase *ψ* (Fig. 2). This interpretation allows us to describe the mechanism underlying the evolution of the bump graphically (Fig. 2).

**Fig. 2.**
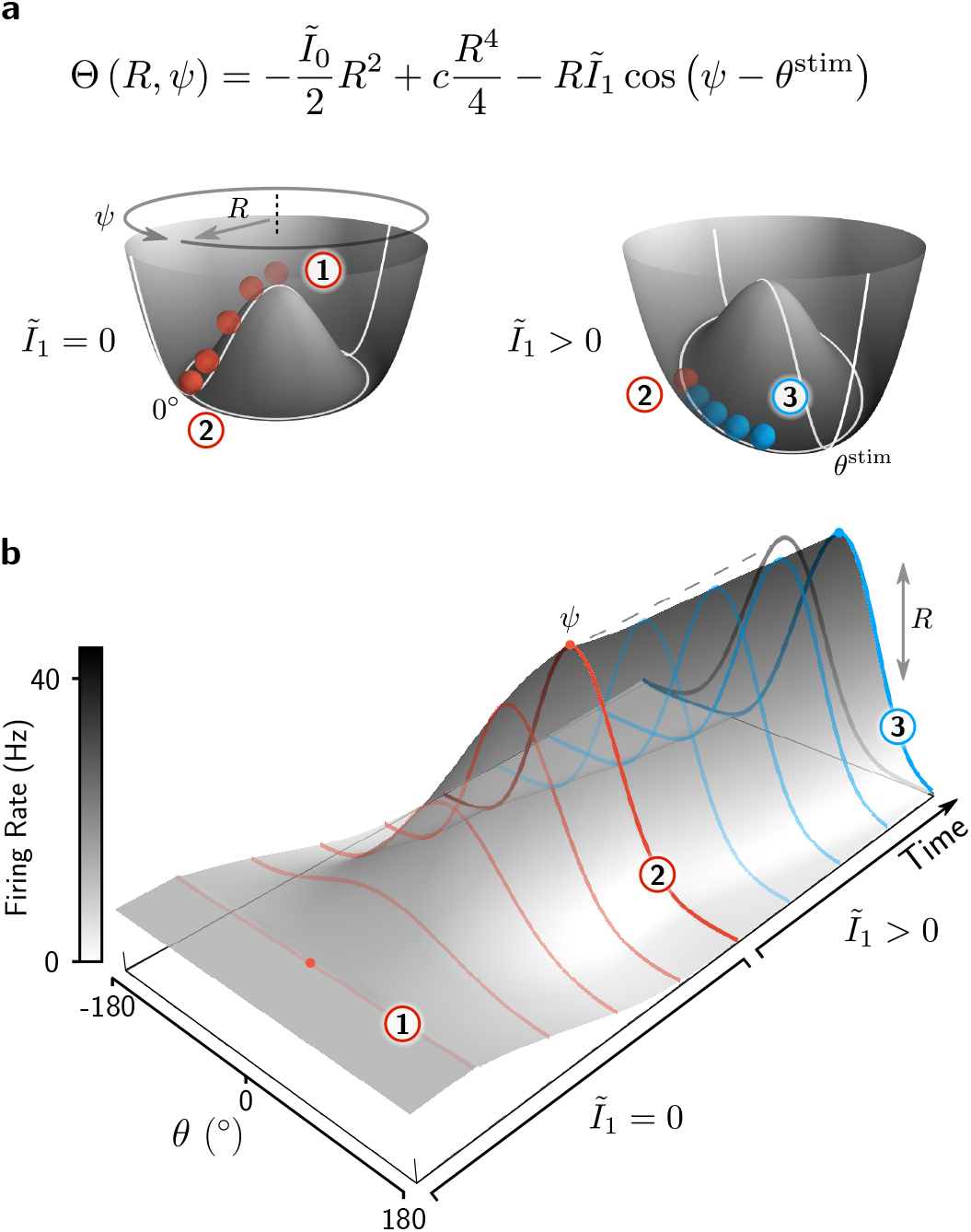
Dynamics of the bump attractor network in the potential landscape. **a**, The equation of the potential corresponding to the dynamics of the bump attractor (Eq. 1) and its geometrical representation. The movement of the red ball in each potential corresponds to the dynamics of bump formation and translation shown in **b**. **b**, Evolution of the network activity from the unstable homogeneous state ① to the stable bump state ② in the absence of external stimuli 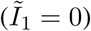, followed by a translation of the bump (transition from ② to ③) towards the location of the stimulus *θ*^stim^ when the external stimulus is present 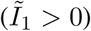. Note that for 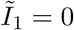, the bump forms at an arbitrary angle *ψ*, determined by noise fluctuations.

In the absence of stimulus input (*I*_1_ = 0), the potential Θ (*R, ψ*) resembles a juice-squeezer-shaped surface with an unstable fixed point at the center, corresponding to a bump with zero amplitude (*R* = 0), i.e. the unstable spatially homogeneous network state (Fig. 2; ①). The circular well of the manifold represents bump attractor states (*R* > 0) that are neutrally stable along the angular dimension *ψ*. The formation of the bump is described by the movement of the particle that, starting at the center of the juice squeezer, rolls downwards towards the stable manifold while increasing its distance to the center, *R* (Fig. 2a, transition from ① to ②). The corresponding evolution of the network activity is shown in Fig. 2b, where the initially homogeneous network activity evolves into a bump state, reaching its steady-state amplitude at point ②.

When presenting a stimulus to the network (*I*_1_ > 0), the radial symmetry of the potential is broken and a deeper region arises at the location of the stimulus *θ*^stim^ (Fig. 2a, right). If the particle has already reached the stable circular manifold at the bottom of the potential, the stimulus will force it to move towards this deeper state along the manifold (Fig. 2a, transition from ② to ③). In terms of the network activity, this corresponds to a movement of the bump towards *θ*^stim^, accompanied by only slight changes in bump amplitude (Fig. 2b). Solving Eq. 1 for small stimulus strengths, we found that the evolution of the phase of the bump depends approximately linearly on time and on the difference between the bump phase and the stimulus direction (Extended Data Fig. 2 and Methods). In contrast, if the particle has not yet reached the stable manifold, i.e., when a stimulus is applied during the initial bump formation, it will cause both a change in bump phase and, together with the internal dynamics, a change in bump amplitude.

In general, successive stimulus frames of a time-varying stimulus drag the bump towards their respective orientations in a continuous fashion (Fig. 1c), with an angular displacement that depends on the current bump amplitude (Extended Data Fig. 1). In sum, we can distinguish three distinct stages of the evolution of the bump during which it becomes increasingly difficult for the stimulus to impact the phase of the bump. Initially, starting from homogeneous network activity, the stimulus orientation strongly impacts the phase of the emerging bump. Second, the bump grows in amplitude, driven by the internal dynamics and the stimulus, and thus successive stimulus frames become less and less effective in moving the bump. Third, when the bump amplitude reaches its steady state, it cannot grow further and the impact of subsequent stimuli becomes constant over time. Equipped with these theoretical results, we can now study different stimulus integration regimes in the network.

### The ring model shows a variety of temporal evidence integration regimes

Our theoretical results showed that the changes in bump phase caused by a time-varying stimulus critically depend on the bump amplitude, suggesting its key role in the integration of sensory stimuli. To investigate this, we ran simulations in which networks with different steady-state bump amplitudes had to estimate the average direction of a sequence of oriented frames, distributed between −90° and +90° as in Fig. 1c. To quantify the dynamics of evidence integration, we used the psychophysical kernel (PK) which measures the influence of sensory evidence during the course of the stimulus (see Methods).

We first considered the case where a trial starts with an initial bump (i.e., the particle is at a position on the circular attractor well of the potential; point ② in Fig. 2), whose amplitude we set by varying the global excitatory drive to the network *I*_0_ and keeping all other parameters fixed. Spontaneous bump states have been observed in prefrontal and visual cortex^38–40^, and we show further below how they can give rise to biases in the estimation process (Fig. 5). The PKs we obtained for different values of *I*_0_ are qualitatively similar (Fig. 3a): they all rise, indicating that the stimulus frames later in the trial have more impact on the final estimate (i.e., the bump phase ψ at the end of the trial) than early stimulus frames. This commonly called “recency” effect becomes weaker as *I*_0_ increases, yielding a decrease of the PK slope (Fig. 3c, blue line). The reason for the general over-weighting of late stimulus information is not trivial and deserves a closer inspection. Essentially, it occurs because the bump amplitude is fixed and each successive stimulus frame leads to a partly discounting of the previous estimate of the stimulus average (leaky integration).

**Fig. 3.**
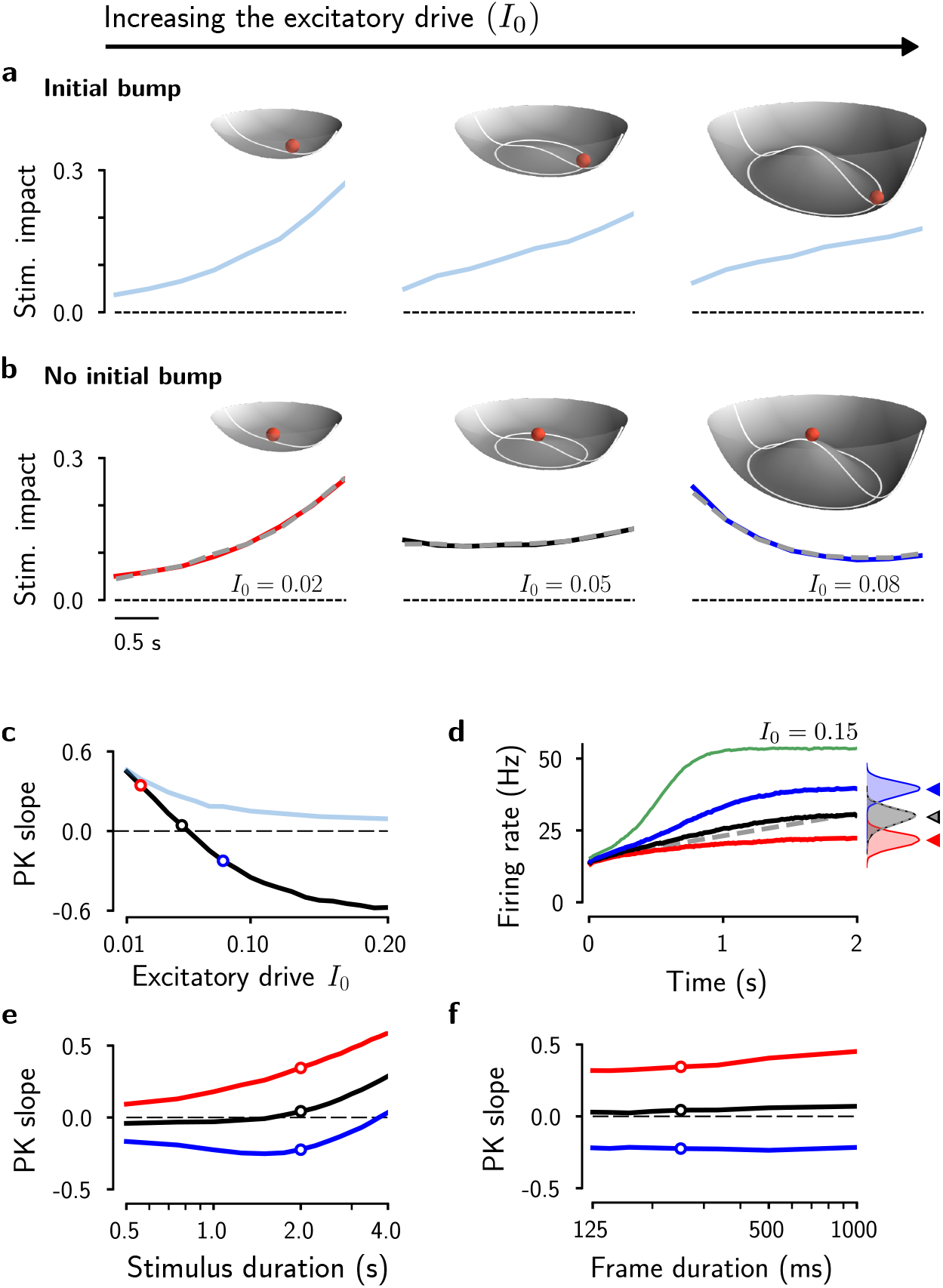
The global excitatory drive I_0_ determines the integration regime of the network. **a**, Psychophysical kernels (PKs) for increasing values of *I*_0_ (bottom, from left to right, *I*_0_ = 0.02, 0.05 and 0.08) with a fully formed bump as the initial condition. The PKs describe the impact of each stimulus frame on the estimated average orientation. Top: For small values of *I*_0_ the potential takes the form of a steep paraboloid where the bump has little space to grow (top left). Intermediate values of *I*_0_ transform the potential into a wide surface similar to a dish with a relative flat central area (top middle). Increasing *I*_0_ widens the potential and gives rise to an increasingly deeper circular manifold at the bottom of the juice squeezer (top right). **b**, PKs as in **a** but with spatially homogeneous initial conditions (flat network activity; particle at the center of the potential). PKs were obtained from numerical simulations of the ring attractor network (colored lines) and from simulating the amplitude equation (dashed gray lines). **c**, PK slope as a function of the excitatory drive *I*_0_, for the spatially homogeneous (black line) and inhomogeneous (blue line) initial conditions. **d**, Evolution of the bump amplitude for different values of *I*_0_. Colored lines correspond to values of *I*_0_ used in **b**. The evolution of the vector length in the PVI is shown for comparison (dotted gray line). Inset: distribution of the bump amplitude at the end of the trial. The histograms for *I*_0_ = 0.05 and for the PVI (in black) perfectly overlap. **e**, PK slope as a function of the stimulus duration, with stimuli composed of frames with a fixed duration of 250 ms. PKs are shown in Extended Data Fig. 5a. **f**, PK slope for a fixed stimulus duration of *T*_stim_ = 2 s as a function of the duration of a stimulus frame. The number of frames ranges from 2 (frame duration 1000 ms) to 16 (frame duration 125 ms).

To fully understand the recency effect, it is necessary to consider how the running average of the stimulus direction can be tracked optimally. Mathematically, we want to keep track of the circular mean of the time-varying stimulus direction *θ*^stim^(*t*), which requires computing the vector sum of the stimulus vectors, each defined by their strength *I*_1_ and their direction *θ*^stim^ (Extended Data Fig. 3a,c; Methods). To compare this optimal solution to the bump attractor network, we derived a dynamical system that continuously tracks the cumulative circular average, the *perfect vector integrator (PVI).* The corresponding two-dimensional equation (Eq. 22 in Methods) turned out to be nearly identical to the amplitude equation but without the intrinsic dynamics of the bump amplitude (the two terms depending on *R* in Eq. 1a). This remarkable equivalence indicates that any deviation from uniform evidence weighting (as in the PVI) must be due to the non-linear amplitude dynamics of the bump attractor model. To gain an intuition on how the amplitude dynamics affects evidence integration, we can again consider the movement of a particle in a potential which for the PVI takes the form of a plane going through the origin and tilted towards *θ*^stim^

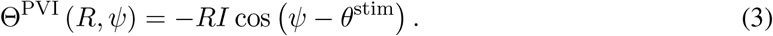

In response to each stimulus the particle moves in both the radial *R* direction and the angular *ψ* direction, thus integrating stimulus vectors. How *R* evolves over time in the PVI depends on the particular combination of the directions of stimulus frames in each trial, but, on average, *R* will grow over time because of a net movement in the direction of the mean of the stimulus distribution 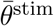 (Extended Data Fig. 3b,d show how the growth rate depends on the width of the stimulus distribution). Because *R* is growing, the angular updates become smaller with each stimulus frame (in other words, as the number of samples grows, the running average direction increasingly relies on the already accumulated information). This is in contrast to the integration process in the bump attractor network with a formed bump in which the bump amplitude is fixed (determined by the shape of the potential; Fig. 2a) and integration occurs only in the angular direction. Because of this, the network gives equal weight to the angular updates over time, thus partly erasing the accumulated stimulus information while over-weighting late stimulus information. The strength of the recency effect depends on the bump amplitude. As described in the previous section, the smaller is *I*_0_ and thus the bump amplitude, the larger is the angular displacement for a given stimulus strength *I*_1_ and the faster the bump can track the orientation of incoming stimuli, resulting in a PK with increasing recency effect (Fig. 3a,c; Extended Data Fig. 1).

Strikingly, when we ran simulations in which the network had to estimate the average stimulus direction starting the trials in the homogeneous network state, with no bump formed (i.e., the particle is at the unstable center position on the potential; point ① in Fig. 2), we observed three qualitatively distinct integration regimes depending on the global excitatory drive to the network *I*_0_ (Fig. 3b; Supplementary Videos 1 to 3). In the first regime, a small *I*_0_ yields again a recency PK (Fig. 3b, left). Here, the steady-state bump amplitude is relatively small, and the bump can only grow very slightly over time (Fig. 3d, red line). Thus, stimulus integration is limited by the small bump amplitude and the network behaves similarly to the previously considered case with the bump formed from the beginning of the trial and with constant amplitude throughout the trial (Fig. 3a, left).

Second, for large *I*_0_, the networks showed a “primacy” effect, that is, early stimulus information now had a higher impact on the final estimation than later stimulus information (Fig. 3b, right; Supplementary Video 3). This can be understood by considering that in this regime the bump amplitude is small at the beginning of the trial but then grows to a large amplitude (Fig. 3d, blue line). Thus, the bump can quickly follow the stimulus direction early in the trial during the formation of the bump, but while growing it becomes increasingly sluggish and stimulus information becomes comparably less effective in displacing the bump. Compared to the PVI, the bump amplitude grows too quickly because it is driven by the internal dynamics in addition to the stimulus (i.e., the potential is not flat as for the PVI but has a peak in the center; Fig. 3b, right inset). The resulting down-weighting of later stimulus frames only occurs while the bump is growing (transition from ① to ② in Fig. 2). Once the steady-state bump amplitude has been reached (corresponding to the attractor well of the potential), the model starts to overweight incoming evidence and, as a consequence, for very long stimulus durations the model will eventually show a recency PK (Fig. 3e, blue line).

Third, for intermediate values of *I*_0_, primacy and recency effects balance out and we obtained an almost perfectly uniform PK (Fig. 3b, middle; Supplementary Video 1). Thus, in this regime the directions of all stimulus frames had the same impact on the final estimate over the whole trial (PK slope = 0; Fig. 3c). In this case, the potential has only a relatively small peak at the center, approximating the flat potential of the PVI (Fig. 3b, middle inset). The growth rate of the bump amplitude is mainly driven by the stimulus as in the PVI (Extended Data Fig. 3d). The value of *I*_0_ in which the model best approximates the PVI depends on the stimulus strength and the distribution of the stimulus directions (Extended Data Fig. 3e). Approximately perfect integration is only possible as long as the bump amplitude can increase (before the well of the potential is reached), such that for long stimulus durations integration becomes again leaky and we obtain again a recency PK (Fig. 3e, black line). Thus, both primacy and uniform PKs are robustly observed for small and intermediate stimulus durations (Fig. 3e; *T*_stim_ ≤ 2 s) while for long durations the recency regime is recovered. The results in Fig. 3 were obtained with time-varying stimuli that change their orientation every 250 ms (frame duration). The PKs remain qualitatively the same when keeping the stimulus duration constant and presenting a variable number of stimulus frames of different duration (Fig. 3f).

### Accuracy of stimulus estimation and categorization

We next quantified the estimation and choice accuracy in the different temporal integration regimes of the bump attractor model when starting the integration without an initial bump of activity. We computed stimulus estimation curves by reading out the bump phase from the network activity at the end of the trial, averaging the estimates across trials, and plotting them as a function of the actual mean orientation 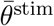 (Fig. 4a). We noticed that independent of the bump amplitude, that was controlled by varying the excitatory drive *I*_0_, the average of the estimates was always close to the true mean direction. We quantified this by computing the estimation bias for 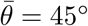 and obtained *b*_est_ (45°) < 1° for all values of *I*_0_. Consistent with this, the slopes of the estimation curve obtained from regression analysis (Methods) were *k*_1_ ≈ 1 in all cases (Fig. 5d, red line). The estimates computed by the ring attractor network were thus approximately unbiased, independent of the temporal weighting of stimulus evidence. The unbiased estimates are a direct consequence of starting the integration process in the neutral center of the potential and the symmetry of the potential (Fig. 2, point ①). The direction of the first stimulus frame determines the initial phase of the forming bump in an unbiased manner. Subsequently, the bump will move, but as long as the directions of the stimulus frames are drawn i.i.d. from some underlying distribution, clockwise and counterclockwise bump displacements are equally likely and the estimate always remains unbiased irrespective of the integration regime.

**Fig. 4.**
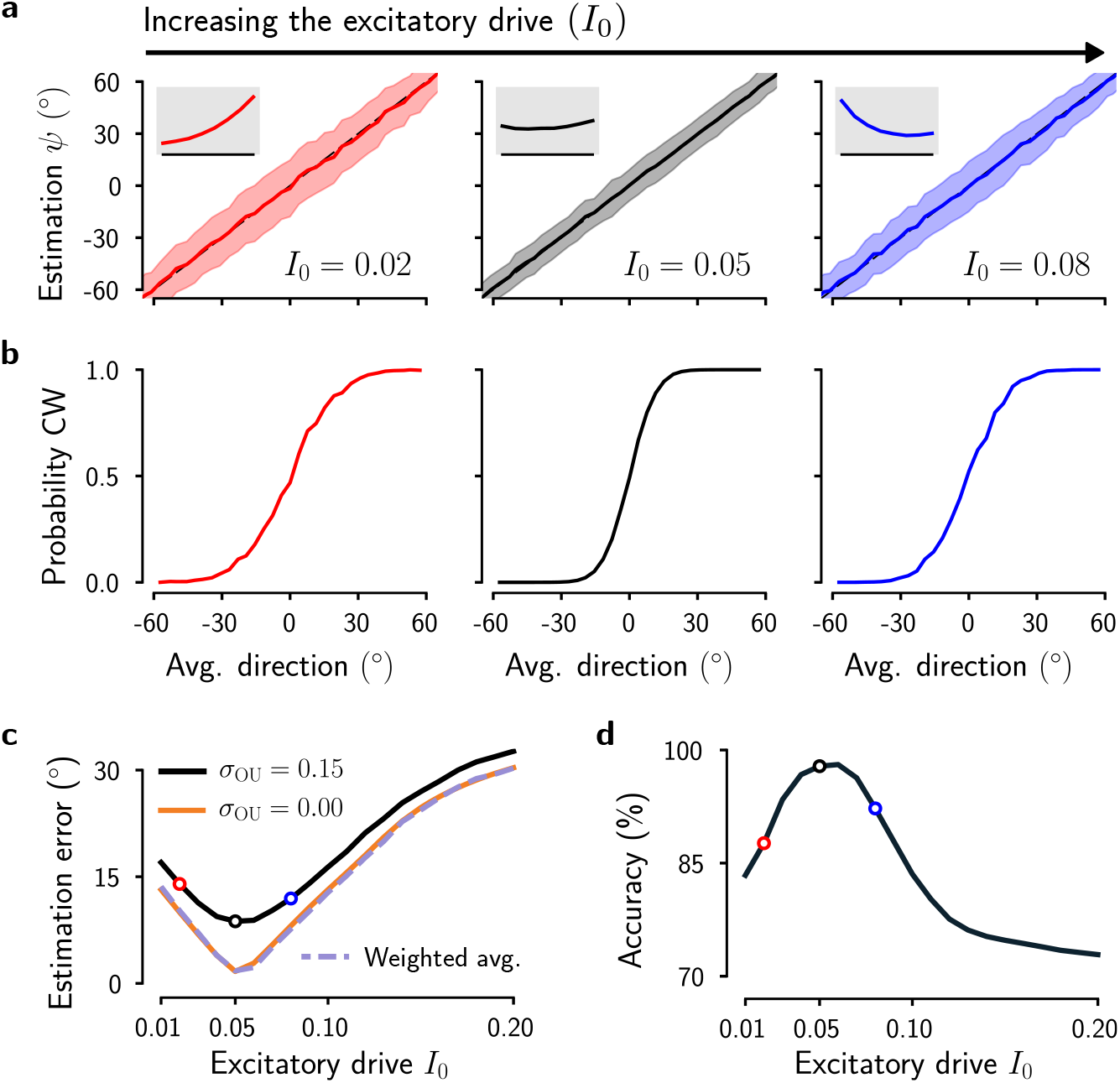
Dependence of estimation and categorization accuracy on the global excitatory drive *I*_0_. **a**, Estimated average direction as a function of the true average of a stimulus composed of 8 oriented frames, for increasing values of *I*_0_ (from left to right, *I*_0_ = 0.02, 0.05 and 0.08). Solid lines indicate the average across trials and the shadings indicate s.d. Insets: corresponding PKs from Fig. 3b. **b**, Probability of a clockwise choice versus average stimulus orientation, for increasing values of *I*_0_. A clockwise/counterclockwise choice corresponds to a positive/negative phase of the bump at the end of the trial. **c**, Root-mean-square error (Methods) of the ring attractor network with noise (*σ*_OU_ = 0.15) and without noise (*σ*_OU_ = 0) as a function of the global excitatory drive *I*_0_. The estimation error for *σ*_OU_ = 0.15 (black line) corresponds to the estimations shown in **a**. The estimation error for a perfect weighted average of the stimulus directions (with the same temporal weighting as the network model but otherwise optimal; Methods) is shown for comparison. Note that the curve corresponding to *σ*_OU_ = 0 has been shifted such that for both conditions the critical value of *I*_exc_ at the bifurcation point is aligned (Methods). **d**, Categorization accuracy as a function of the excitatory drive *I*_0_, for a fixed average stimulus orientation 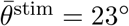.

**Fig. 5.**
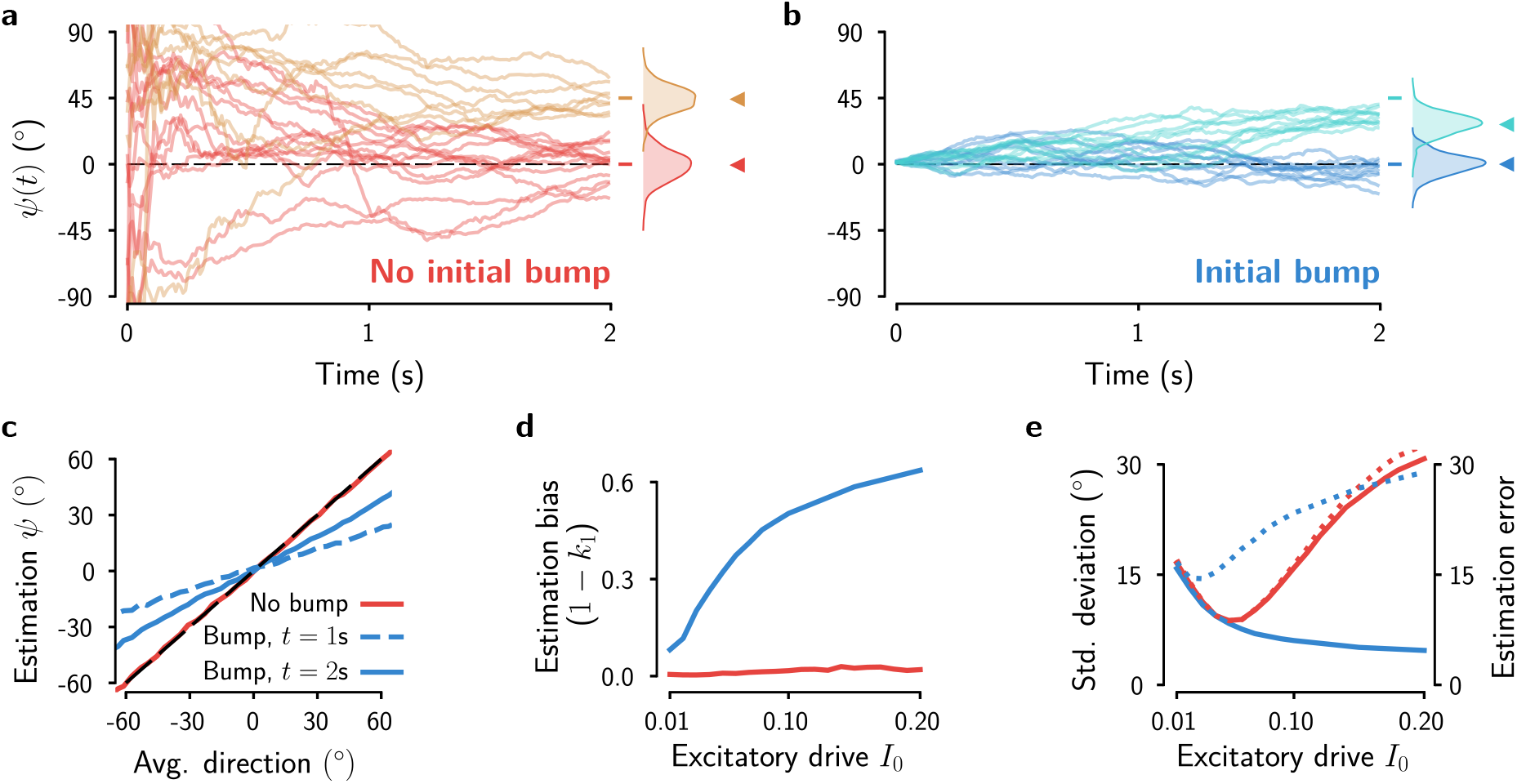
Impact of the initial condition on the bias and variance of the stimulus estimates. **a**, Bump phase in example trials with 0° and 45° average stimulus directions (*N* = 8 stimulus frames, stimulus duration *T*_stim_ = 2 s) when starting the trial with homogeneous network activity (i.e., no bump formed). Inset: distribution of the phase of the bump at the end of the trial. **b**, Bump phase for the same network and the same stimuli as in **a**, but starting the trial with a fully formed bump centered at 0°. **c**, Continuous estimations as a function of the mean stimulus direction 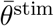 for two different stimulus durations (no bump condition in red, bump condition in blue). For the no-bump condition, the average estimation curve does not depend on stimulus duration (red line) and it overlaps with the diagonal (dashed black line). **d**, Systematic bias of the estimates as a function of the global excitatory drive *I*_0_ to the network (stimulus duration *T*_stim_ = 2 s). The estimation bias is defined as one minus the slope of the estimation curve (*k*_1_ in Eq. 32, Methods). **e**, Standard deviation (solid lines, left axis) and root-mean-square estimation error (dotted lines, right axis) of the final bump position as a function of the global excitatory drive *I*_0_ for the same trials as in **d**.

However, the trial-to-trial variability of the estimates (Fig. 4a, shaded areas and Fig. 4c, black line) depended on the integration regime. Several factors could give rise to estimation errors in the attractor model and may thus contribute to this dependence: (1) internal noise and stimulus fluctuations of magnitude *σ*_OU_ (Methods), (2) sub-optimal evidence integration due to non-uniform evidence integration (i.e., non-uniform PKs), and (3) additional impact of the non-linear internal dynamics on evidence integration not captured by the PK. We first considered the impact of the noise. We ran simulations of the network without noise (*σ*_OU_ = 0) and obtained, as expected, a general decrease of the estimation errors (Fig. 4c, orange line). We next tested to what degree the remaining estimation errors can be accounted for by the non-uniform evidence weighting in the primacy and recency regimes. To show this we compared the dependency of the estimation error on *I*_0_ of the attractor model with the estimation error that was obtained when computing the true average of the stimulus directions with the PVI but weighted with the PKs from the bump attractor model (Fig. 4c, purple dashed line). The obtained estimation errors are almost indistinguishable from the errors of the noise-less bump attractor network (Fig. 4c). Thus, the increase of estimation errors for non-optimal *I*_0_ (corresponding to values of *I*_0_ different from 0.05 in Fig. 4c) can be attributed to the suboptimal, non-uniform, temporal weighting captured by the PK. Because the non-linear internal dynamics of the ring model can only yield an approximately uniform PK, the estimation errors are non-zero even for the best *I*_0_ for a given stimulus distribution, in particular when the stimulus distribution is wide (Extended Data Fig. 3e). Moreover, the model PKs change with the duration of the stimulus and thus the estimation error also shows a time-dependence (Extended Data Fig. 5). Together, these analyses show that the estimation accuracy in the bump attractor network is limited by noise and by the non-linear internal dynamics causing non-uniform evidence integration, whereas further contributions of non-linearities (that are not captured by the PK) are negligible.

A direct consequence of the trial-to-trial variability of the estimates is that they limit the accuracy of a categorical decision that is based on the read-out of the bump phase. This can be seen when computing psychometric curves by interpreting a positive bump phase at the end of the trial as a clockwise choice and a negative bump phase as a counterclockwise choice, respectively (Fig. 4b). As expected from the stimulus estimation results, the psychometric curve for the uniform integration regime had the highest slope (Fig. 4b), and decision accuracy is maximal for the *I*_0_ that gives rise to a uniform PK (Fig. 4d).

### Initial bump states cause an estimation bias

We next investigated in more detail the accuracy of stimulus estimation in the network with an initially formed bump (Fig. 3a), in particular the impact of the initial bump phase on the estimation process. In psychophysical experiments this initial bump position could be determined by a reference line shown before the onset of the stimulus^2,4^ or it could be related to the subject’s prior expectation before the start of the stimulus^5^. Single trials are illustrative of the effect of the different initial conditions (Fig. 5a,b). When starting a trial in the homogeneous network state, the first stimulus frame determines the phase where the bump emerges (Fig. 5a), and the bump tracks an estimate of the average direction as the trial progresses. As described above, the network’s estimates were unbiased in this case (Fig. 5c,d; red line). In contrast, when starting with a fully formed bump at a particular position as the initial condition, the bump phase was biased towards the initial phase of the bump (Fig. 5b,c). As the network integrates the stimulus, the bump moves successively closer to the true average orientation, leading thus to a decrease of the systematic deviations from the true average with stimulus duration (Fig. 5c, blue lines). The magnitude of the estimation bias for a fixed stimulus duration of 2 s increased with the global excitatory drive *I*_0_ to the network (Fig. 5d). The reason is that a higher *I*_0_ yields larger bumps that are more sluggish and thus need more time to overcome the bias caused by the initial bump position. While the variability of the estimates in the biased case is generally lower than in the unbiased case, the overall estimation error (bias + variance) is higher (Fig. 5e). To confirm that the bias is actually caused by the pre-determined phase of the activity bump at the beginning of the trial and not a consequence of starting the trials with a fully formed bump, we ran simulations with a fully formed initial bump with a phase determined by the first stimulus frame and indeed observed no bias (Extended Data Fig. 4).

In sum, starting with an initial bump led to recency PKs (Fig. 3c) and, when setting the initial phase to some pre-determined value it yielded estimates that were biased towards the initial bump phase. It is worth noting that here we considered two extreme cases, either starting in the neutral condition (no initial bump) or with a fully formed initial bump. Intermediate scenarios (e.g. starting with a partially formed bump of lower amplitude or starting with a fully formed bump in only a fraction of trials) would lead to PKs, biases and estimation errors that lie between the extreme cases.

### The ring model explains heterogeneity in integration dynamics in humans

We tested whether human subjects show variations in their integration dynamics in a stimulus averaging task and whether the ring model could parsimoniously account for those variations (Fig. 6). In this category-level averaging task, a visual stream of eight oriented Gabor patters was presented to participants^6^ (Fig. 6a). Each frame had an orientation between −90° and 90° and a fixed duration of 250 ms, conforming a stimulus with total duration of *T*_stim_ = 2 s. At the end of the stream participants reported whether, on average, the tilt of the eight frames fell closer to the cardinal {0°, 90°} or diagonal {−45°, 45°} axes. On average, participants were able to combine the evidence favouring either option improving their accuracy in trials with higher evidence^6,7,41^.

**Fig. 6.**
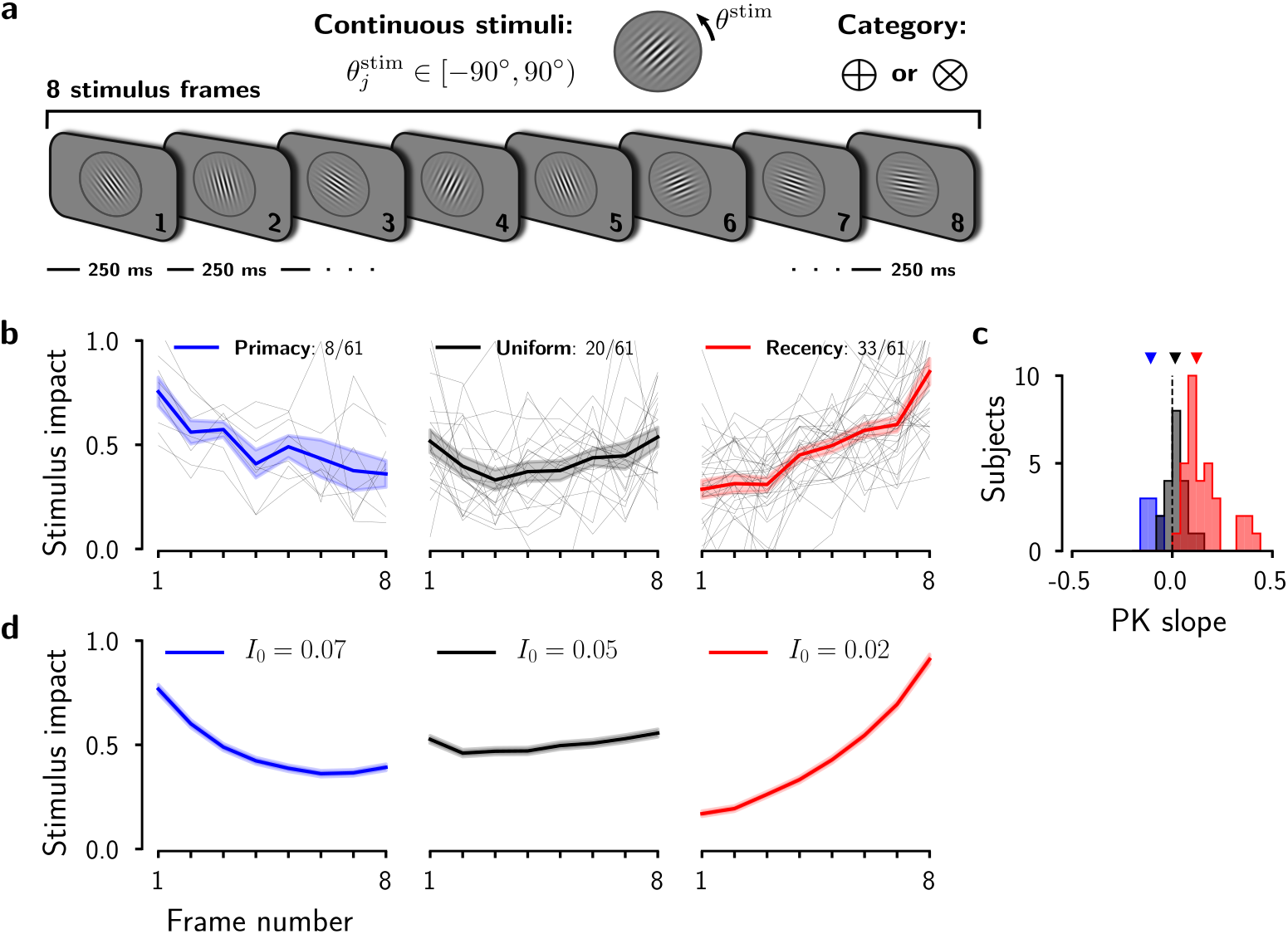
The ring attractor model accounts for experimentally observed psychophysical kernels. **a**, Category-level averaging task^6,7,41^. Participants reported whether, on average, the tilt of eight oriented Gabor patterns fell closer to the cardinal or diagonal axes. **b**, PKs obtained from human subjects were heterogeneous. Thin lines represent the PKs of individual subjects and colored lines the group averages. **c**, Distributions of the slopes of the PKs for each type of integration dynamics. The top triangle indicates the median for each case. **d**, PKs obtained by fitting the ring model to the experimental data by varying the excitatory drive *I*_0_.

We merged the data from three studies^6,7,41^ carried out using very similar experimental conditions (*n* = 61 subjects, Methods). We computed their individual PKs (Methods) and classified the subjects according to their temporal weighting behavior (primacy, uniform, recency). As previously reported^7^, across participants there was a tendency for late temporal weighting (recency). However, subject-by-subject analysis revealed a broad range of integration dynamics (Fig. 6b,c; individual PKs are shown in Extended Data Fig. 6). The majority of participants weighted more the sensory evidence during the late periods of the stimulus (33 out of 61 participants), yet a substantial proportion of the participants weighted the evidence approximately uniformly (20 out of 61 participants). Finally, a minority of participants (8 out of 61 participants) weighted early frames of the stimulus more that late frames, showing a primacy PK. Qualitatively, the ring model could capture these different temporal integration dynamics observed in the psychophysical experiments (Fig. 3). To assess whether the model could explain the data in a quantitative manner, we performed simulations with the same stimulus statistics as in the behavioral experiment. We systematically adjusted both the global excitatory drive to the network, *I*_0_, and the noise level, *σ*_OU_, such that the ring network reproduced, on average, the experimental results (Fig. 6b,c), characterized by the slope of the across-subjects PKs (Fig. 6b) and the average performance (not shown) for each integration regime (primacy, uniform and recency). The inter-subject heterogeneity in temporal evidence weighting could thus parsimoniously be explained by varying the the overall excitatory drive that determines the amplitude of the bump, while adjusting the noise fluctuations to match the performance of subjects.

It is conceivable that the global excitatory drive does not only differ between individuals but that it could also be modulated on a trial-to-trial basis, such that it may therefore change the time-course of evidence integration depending on the current task conditions. Such a task-driven change has previously been observed in a similar task as in Fig. 6 that studied the mechanisms of perceptual choices under focused and divided attention^41^. In that task, two spatially separated stimulus streams were presented simultaneously and subjects had to integrate either a single or both streams in a trail-to-trial basis. We built a network model consisting of two ring circuits (Methods) and found that the results of this experiment can be explained by a change in the excitatory drive that controlled the allocation of a fixed amount of resources between the two circuits (Extended Data Fig. 7). In sum, our model suggests that the overall excitatory drive determines the integration regime. Control signals like top-down attention signals or neuromodulatory gain changes can impact the dynamics of evidence integration and the model makes specific predictions about the underlying changes in neural activity.

### Bump attractor dynamics links integration dynamics and reaction times

Finally, to derive specific predictions from the model that go beyond the shape of the PK, we extended the model by coupling the ring integration circuit to a canonical decision circuit and studied the decision build-up in the different evidence integration regimes (Fig. 7). In this two-circuit model (Fig. 7a; Methods), evidence about the average stimulus direction is integrated in the phase of the activity bump in the integration circuit and then converted to a categorical choice in the decision circuit. The decision build-up is triggered by an urgency (or choice commitment) signal at the offset of the fixed-duration stimulus.

**Fig. 7.**
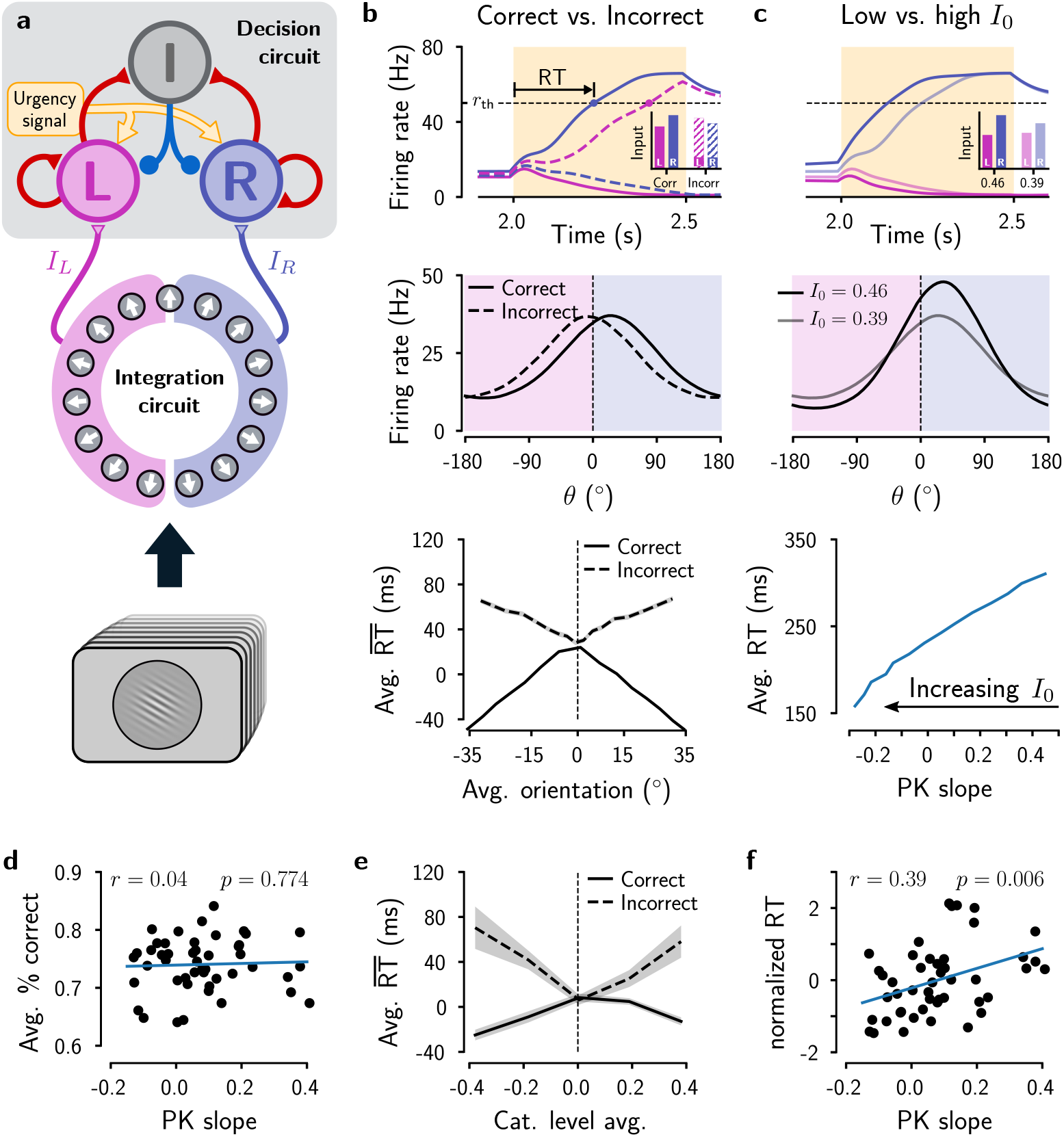
Dependence of reaction times on evidence strength and on the shape of the psychophysical kernel. **a**, Ring integration circuit coupled to a categorical readout circuit (decision circuit; Methods). Left (**L**) and right (**R**) excitatory populations in the decision circuit receive inputs, *I_L_* and *I_R_*, from their corresponding side of the ring. The inhibitory population (**I**) promotes winner-take-all competition^8^. The decision process is prompted by the activation of an urgency signal at the end of the stimulus presentation that initiates the competition. **b**, Dependence of reaction times (RTs) on choice outcome and stimulus evidence in the model. Top, average activities of the **L** and **R** populations for correct (solid lines) and incorrect (dashed lines) trials for stimuli with average direction of 15°. RT is measured from the onset of the urgency signal (at *t* = 2 s; shaded area) until one of the firing rates reaches the threshold *r*_th_ = 50 Hz. The inset shows the summed bottom-up inputs, *I_L_* and *I_R_*. Middle, average activity of the integration circuit at the end of the stimulus presentation. Bottom, Dependence of RTs on stimulus evidence for correct and incorrect trials. **c**, Dependence of RT on the excitatory drive *I*_0_ in the model. Top and middle panels as in **b** but for two different levels of the excitatory drive and correct trials only. Bottom, Average RTs as a function of the PK slope. **d-f,** Performance and RTs in human subjects (*n* = 47). **d**, The percentage of correct trials is not correlated with the PK slope (*r* = 0.04, *P* = 0.774, *t*-test). **e**, Average RTs as a function of the category level average for correct and incorrect trials. RTs for each condition were computed relative to the trial-averaged RT across all trials, 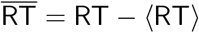. Errorbars indicate the SEM across subjects. **f**, Subject’s RTs (z-scored values for each data subset, Methods) vs. their PK slopes show a significant correlation (*r* = 0.39, *P* = 0.006, *t*-test).

We observed that the reaction time (RT), from stimulus offset to reaching a categorical choice, depends on the trial outcome (correct or incorrect) and on the accumulated stimulus evidence (Fig. 7b, bottom). RTs for correct trials are shorter and they decrease with stimulus strength, whereas RTs for incorrect trials are longer and they increase with stimulus strength. This can be understood by considering how the decision build-up in the decision circuit depends on the position of the activity bump in the integration circuit. For a given average stimulus direction, the activity bump moves further away from the decision boundary in correct trials than in incorrect trials (Fig. 7b, middle). Thus, one of excitatory decision populations (**L** or **R**) receives a stronger input signal than the other population and, due to winner-take-all competition, its firing rate rapidly increases, reaching the decision threshold quickly (Fig. 7b, top). In incorrect trials, the resulting input to the **L** and **R** populations is more similar and the decision build-up takes longer. This effect gets more pronounced for larger average orientations (i.e., the bump moves further away from the decision boundary in correct trials and less so in incorrect trials). We found that both signatures, the decrease of RTs with accumulated stimulus evidence for correct trials and the increase of RTs for incorrect trials, are also present in the experimental data obtained from human subjects (Fig. 7e).

Moreover, the model predicts a direct relationship of RTs and the shape of the psychophysical kernel (Fig. 7c). Different psychophysical kernels in the model are obtained by varying the excitatory drive to the network (Fig. 6). Increasing the excitatory drive leads to an increase of the bump amplitude and a transition from recency over uniform to primacy weighting (Fig. 3b,c). The bump amplitude, in turn, affects the RT in the two-circuit model (Fig. 7c). RTs are larger for small bump amplitudes because the decision circuit receives weaker input and takes therefore longer to reach the decision threshold. Taken together, the model predicts that primacy temporal weighting should be accompanied by shorter RTs and recency weighting by longer RTs, i.e., RT should increase with the slope of the PK (Fig. 7c, bottom). We tested this non-trivial prediction in the experimental data and indeed found a significant correlation between the subject’s RTs and PK slopes (Fig. 7f; *r* = 0.39, *P* = 0.006, *n* = 47, *t*-test). It is unlikely that the differences in RTs could be explained by a different fraction of correct trials depending on the PK slope because we did not find a correlation between PK slope and the fraction of correct trials (Fig. 7d; *r* = 0.04, *P* = 0.774, *n* = 47, *t*-test). In sum, the two-circuit model provides a mechanistic link between neural population activity, evidence integration dynamics and reaction times.

## Discussion

We investigated how the classical continuous ring attractor model^28,30,33^ integrates a time-varying stimulus in the amplitude and the phase of the network’s activity bump. We found that the model can approximately compute the time integral of stimulus vectors defined by the strength and the direction of the stimulus. This operation can be understood in terms of a population vector^42^. But while a population vector is typically used to read out direction information from a neural population (by weighting the activity of individual neurons according to their preferred direction), the population vector encoded in the amplitude and the phase of the activity bump results from a temporal integration process and is continuously updated in response to incoming stimulus evidence.

We studied the neural network mechanism underlying this integration process and identified how the model integrates a sequence of stimulus frames of varying direction and constant strength, e.g. a sequence of visual gratings with different orientation but the same contrast. Analytically and through simulations we confirmed that the phase of the activity bump tracks the running circular average of the stimulus orientations (for variable stimulus strengths we would get a weighted circular average). Concurrently, the amplitude of the bump evolves depending on the dispersion of the stimulus directions. The dynamical regime in which the model closely approximates perfect vector integration depends on the stimulus statistics as well as the stimulus strength and duration (Extended Data Fig. 3e; Extended Data Fig. 5b). It is characterized by a relatively wide and shallow potential so that the dynamics of the bump is dominated by the stimulus and not by the non-linear internal dynamics (Fig. 3). The limiting factor imposed by the internal dynamics of the network is that the bump amplitude cannot grow beyond a maximum value that depends on the model parameters and is proportional to the overall excitatory drive to the network. When the maximum bump amplitude is reached, which will eventually happen for long stimulus durations, the model has reached the neutrally stable ring attractor and can still track the average stimulus phase. However, the model is then over-weighting later stimulus frames, similar to a leaky integrator. In addition to this recency effect, the model can also show primacy temporal weighting when the internal network dynamics contributes to a rapid initial growth of the bump and initial stimulus frames have a relatively higher impact on the evolution of the bump phase. Independent of the temporal weighting regime, the average orientation estimates of the model were always unbiased; the variance of the estimates is however minimal when the model acts as perfect vector integrator and increases for non-uniform temporal weighting (Fig. 4). In sum, we have uncovered the fundamental mechanism underlying stimulus integration in the phase and the amplitude of the bump of the ring attractor model and the corresponding computation of the running circular mean. Our results suggest ring attractor dynamics as a potential neural mechanism for the estimation of average circular stimulus features and for perceptual discriminations based on the same features.

### The role of the global excitatory drive

The key parameter that controls the integration dynamics in the bump attractor model is the global excitatory drive to all neurons in the network, *I*_0_ (Fig. 3). Keeping all other parameters fixed, *I*_0_ determines the depth and the width of the potential (Extended Data Fig. 1) and thus the maximal bump amplitude and the intrinsic network dynamics. We showed that by varying *I*_0_, the model could account for the heterogeneity in how human observers weighted sensory evidence across a stream of oriented stimulus frames (Fig. 6) as well as for their reaction times (Fig. 7). In addition to a different level of global excitatory drive, subjects are likely to show other differences as well, for example a different E/I ratio or a different gain of the neural circuits involved in evidence integration. Effectively, in the model these inter-subject differences can be captured by different values of *I*_0_. In particular, it can be shown that changing the slope of the neuronal transfer function is mathematically equivalent to a change in *I*_0_. The bump attractor model can also account for inter-subject differences in evidence weighting by a less parsimonious explanation that involves the joint modulation of the global excitatory drive, the internal noise and the initial state of the network. That is, constraining *I*_0_ to a narrow range while modulating the initial bump amplitude or the fraction of trials that start with an initial bump would also result in different psychophysical kernels (Fig. 3c). Initial bump states generally yield more recency and larger estimation errors (Extended Data Fig. 4e,f). However, our experimental data does not support this hypothesis because the accuracy of human subjects did not depend on the slope of the PK (Fig. 7d).

The global cortical gain can also be modulated in a task-dependent manner across time in individual subjects, for example controlled by top-down signals and neuromodulators such as noradrenalin originating from the locus coeruleus^43–45^. We showed that this can explain attention-driven differences in evidence integration (Extended Data Fig. 7). Moreover, a time-varying neural gain or excitatory drive in the model can switch the network from an integration regime into a working memory regime. During stimulus estimation, the excitatory drive should be moderate so that the network would have a relatively shallow potential and would act as a perfect vector integrator. Subsequently, after stimulus presentation, an increase of *I*_0_ would make the potential wider and deeper and thus increase the stability of the bump against noise and against further incoming stimuli as is needed for distractor-resistant working memory^30,46^. This transition could also be gradual, for example caused by a ramping *I*_0_ ^47^.

### Mechanisms underlying biases in perceptual estimation and categorization tasks

Starting the integration process in the model with a formed activity bump shifts temporal integration towards recency and can introduce estimation biases towards the initial bump position (Figs. 3 and 5). The existence of bump states before stimulus presentation is supported by neural population recordings^38–40^. A recent paper^5^ investigated how prior expectations influence motion direction estimation in humans, and found attractive biases towards the predictive cued direction, and a correlation between reported directions and directions decoded from MEG activity that emerged before stimulus onset. These findings are consistent with the initial bump condition in our model (Fig. 5).

Moreover, it has been shown in combined discrimination and estimation tasks that stimulus estimation is influenced by a categorical decision causing post-decision biases^2,4,20,21^. One hypothesis is that these bias effects are mediated through attention signals and through changes in global gain^4,21^. In the bump attractor model, attention related signals could be modeled either as global changes on the excitatory drive to the neurons or as spatially modulated signals targeting specific regions of the network. For example, in the seminal work of^2^ subjects showed a repulsive bias of direction estimates (away from a reference) while performing a fine discrimination-estimation task in which they first had to report whether a pattern of moving dots was clockwise or counterclockwise of a reference. In contrast, subjects showed an attractive bias (towards a reference) in a similar coarse discrimination task in which they had to report whether the dots moved towards or 180° away from the reference. In the model, a spatially modulated attention signal, targeting the location of the two possible choices in the ring, could potentially reproduce the experimentally observed biases. During the fine discrimination task, the attention signal would attract the bump towards the clockwise or counterclockwise directions away from the reference causing a repulsive bias, while the same mechanism would cause the attraction bias during the coarse discrimination task.

Overall, our network model provides a comprehensive computational framework for investigating the neural mechanisms underlying stimulus estimation and perceptual categorization and their interaction in future studies.

### Experimental predictions provided by our model

Our model accounts for experimentally observed PKs (Fig. 6) and we have confirmed the predicted relationship between PKs and reaction times (Fig. 7) in human subjects. Moreover, the bump attractor model yields several predictions that can be tested in human or monkey experiments to validate our theory.

First, our model predicts characteristic changes of the evidence integration dynamics when changing the stimulus properties (e.g. its strength or duration), or its statistics. PKs should shift towards recency for longer stimulus durations (Fig. 3e), which could be tested in a task with randomly interleaved trials of different durations. A similar shift should be observed for stronger, i.e., high contrast stimuli. Additionally, the model predicts changes for manipulations of the generative distribution of the stimuli. For example, for a subject with a particular *I*_0_, broader distributions of the stimulus directions would shift the PK towards primacy and also increase the estimation error (Extended Data Fig. 3b,e). Alternatively, subjects could also internally adapt their *I*_0_ while keeping their PK constant to minimize the error (Extended Data Fig. 3e). Overall, these predictions are constrained by the fundamental mechanisms that govern the integration dynamics in the bump attractor network which are fully captured by the amplitude equation (Eq. 1).

Second, the dynamics of evidence integration crucially depends on the global excitatory drive or gain of the model (Fig. 3) and this could be tested experimentally using pharmacological or optogenetic manipulations. A recent study has compared stimulus integration in human participants in sessions where they have been administered the NMDA receptor antagonist ketamine or a placebo^48^. Reduced excitability under ketamine led to more recency PKs (more leaky integration), as predicted by a reduction in excitatory drive in our model.

Third, the central prediction of our model is a systematic relationship between the amplitude of the population response and the rate of change of its phase (Extended Data Fig. 1). This could be validated in simultaneous multi-unit recordings in monkeys performing an orientation averaging task. It would require decoding the current estimate of the average direction during evidence integration and measure the impact of individual stimulus frames on this estimate. The change of phase in response to the same orientation difference (between stimulus frame and current average) should decrease in a manner inversely proportional to the population firing rate. Thanks to recent advances in fine-grained decoding of sensory and decision information across multiply brain areas from MEG activity^49^, it may be possible to test the same prediction in humans.

### Comparison with other models

The investigation of neural networks that can integrate information across time has a long history^50^. Neural integrators are usually categorized into either rate based^51,52^, accumulating the signal in the graded firing rate of the neurons, or location based, in which the integrated information is encoded in the localized activity of the neurons, for example tracking the head direction in the phase of an activity bump^33,34,53^. The network model presented in this work belongs to the latter category and its activity is governed by bump attractor dynamics as in models of the head direction system. However, the underlying integration process is different. Head-direction models integrate self-motion information that is presented in the form of angular head velocity signals, while visual cues only serve as calibration to partially reset the network. In contrast, in our model the basic operation involves integrating a time-varying sensory stimulus to compute its average direction. Crucially, we have shown that this computation is mediated by the transient interplay between the location of the bump and also its firing rate amplitude, a feature that, to the best of our knowledge, has remained unexplored until now.

One previous study investigated whether a similar continuous recurrent network could identify stimulus features as analog quantities, and also discriminate among a set of discrete alternatives^29^. By changing the underlying dynamical regime, their network was able to perform both computations in two separate tasks. In a direction identification task, they investigated how an orientated stimulus represented in an activity bump is affected by perturbations mimicking cortical microstimulation experiments, similar to the effect of distractors in working memory models^30^. They found that two competing localized bumps can emerge, encoding both the visual stimulus and the microstimulation, and elicit either a stochastic winner-take-all or a spatial averaging process depending on the distance between the two bumps. Overall their integration mechanism is different from the temporal integration studied here, which relies on the dynamics of a single activity bump in response to a time-varying stimulus that achieves the computation of the running average orientation.

Mathematically, the bump attractor model can be viewed as a generalization of the discrete attractor model of perceptual decision making^8,9,11,54^. However, the dynamics of evidence integration in the two models are fundamentally different. Discrete attractor models have a double-well potential that usually leads to a primacy temporal weighting because once the system settles into one of the two attractors it remains there until the end of the trial. Recently we have described how uniform and recency weighting can be obtained when fluctuations in the stimulus together with the internal noise are strong enough to overcome the categorization dynamics and cause transitions between the attractor states^54^. In contrast to this, in the bump attractor model a continuous integration process without abrupt transitions occurs in the neutrally stable ring manifold and the stimulus always impacts the bump phase, yielding recency without the need for strong fluctuations (Fig. 3a).

### Low-dimensional ring attractor dynamics in neuronal circuits

Is bump attractor dynamics realized in a cortical circuit a plausible neural mechanism for stimulus estimation? A bump attractor realized in a physical ring layout has recently been found in the fly’s heading direction system^36,55,56^. Taking advantage of simultaneous recordings of multiple neurons it has been confirmed that the high-dimensional neural population activity in the mammalian head direction system evolves along a one-dimensional ring attractor manifold^35^ consistent with the dynamics of ring models^32–34^. We hypothesize that a similar topological ring manifold may be embedded in the neural population activity of cortical areas that are involved in stimulus estimation. In order to test experimentally whether bump attractor dynamics underlies orientation averaging, it would be interesting to record neural activity from parietal and frontal areas of non-human primates that perform an estimation task and apply manifold identification approach as in Chaudhuri et al.^35^. Such data would also allow for testing whether trial-to-trial variability of neural firing rates and the estimates reported by the subject are related as predicted by the bump attractor model, as in prefrontal cortical neurons during spatial working memory ^31^. In fact, our work suggests that the same neural circuits that are involved in working memory may also be able to carry out stimulus averaging in perceptual estimation tasks. Bump attractor dynamics may thus be a unifying neural mechanism underlying both working memory and evidence integration over prolonged timescales.

## Methods

### Ring model

The dynamics of the bump attractor model are described in terms of the effective firing rate, *r* (*θ, t*), of a neural population arranged in a ring, *θ* ∈ [–*π, π*) (Fig. 1a), obeying the integro-differential equation^27,28,57^

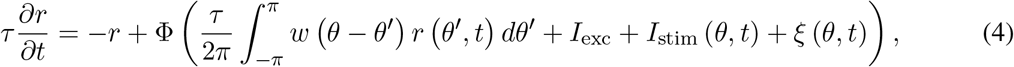

where *τ* is the neural time constant and Φ(·) is the current-to-rate transfer function. The synaptic input to a neuron with preferred orientation *θ* consists of a recurrent current due to the presynaptic activity at a location *θ*′ with a weight *w*(*θ* – *θ′*), and external currents *I*_exc_ + *I*_stim_ (*θ, t*) plus fluctuations *ξ*(*θ, t*). The connectivity profile *w*(*θ*), is written in terms of its Fourier coefficients, *w_k_*(*k* = 0,1, 2,…) and represents the effective excitatory/inhibitory coupling, and can therefore include both positive and negative interactions. The inset in Fig. 1a shows an example of Mexican-hat type connectivity with strong recurrent excitation and broad inhibition. External inputs are divided into a global net excitatory drive *I*_exc_ that modulates the excitability of the network and a time-varying input *I*_stim_ that represents the sensory stimulus. In general, the sensory stimulus can be written in terms of its Fourier coefficients *I_k_, k* = 1, 2,… with directions *θ_k_*(*t*). Throughout this work we will focus on an input of the form *I*_stim_(*θ, t*) = *I*_1_ cos (*θ* – *θ*^stim^(*t*)) to model a stimulus with constant strength *I*_1_ and a time-varying orientation *θ*^stim^(*t*) but our derivations are equally valid for time-varying stimulus strengths *I*_1_. A detailed description of the stimuli used in the simulations is given below (Model simulations). Finally, the fluctuation term *ξ*(*θ, t*) reflects the joint effect of the internal stochasticity of the network and temporal variations in the stimulus. For simplicity, we model these fluctuations as independent Ornstein-Uhlenbeck processes for each neuron, with amplitude *σ*_OU_ = 0.15 and time constant *τ*_ou_ = 1 ms. Tab. 1 summarizes the values of the model parameters used in this work.

**Table 1.** List of the parameters of the ring network and the stimulus.

| **Ring network (Eq. 4)** | *Symbol* | Value |
| --- | --- | --- |
| Time constant of the network | $\tau$ | 20 ms |
| Fourier modes of the connectivity | $w_k, k = 0, 1, 2$ | -2, 1, 0.5 |
| Global net excitatory drive | $I_{\text{exc}}$ | $\in [1.01, 1.2]$ |
| Noise term | $\xi(\theta, t)$ | Eq. 4 |
| Noise amplitude | $\sigma_{\text{OU}}$ | 0.15 |
| Noise time constant | $\tau_{\text{OU}}$ | 1 ms |
| Transfer function | $\Phi(\cdot)$ | Eq. 5 |
| Critical excitatory drive | $I_{\text{crit}}$ | 1 |
| Stationary FR at $I_{\text{exc}} = I_{\text{crit}}$ | $r_{\infty}$ | 12.5 Hz |
| **Stimulus (Eqs. 27 and 28)** | *Symbol* | Value |
| Stimulus amplitude | $I_1$ | $5 \cdot 10^{-3}$ |
| Duration of the stimulus | $T$ | 2 s |
| Number of stimulus frames | $N$ | 8 |

The effective network, Eq. 4, assumes that excitatory and inhibitory neurons follow the same dynamics, and thus can be grouped together, greatly simplifying the mathematical analysis. Nevertheless, a mathematical analysis of the more general model with separated pools of excitatory and inhibitory neurons is possible and is expected to give the same results^58–61^.

#### The bump attractor

In the following we describe the conditions for which the ring network shows a stable bump of activity. We consider the quadratic/square root transfer function^34,62^

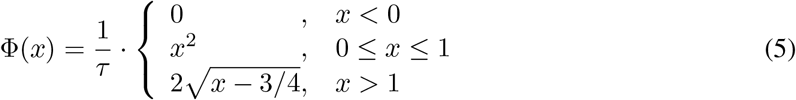

where *x* is the total synaptic input arriving to the neurons. Other continuous non-linear transfer functions will lead to the same qualitative behavior provided they are a monotonically increasing function of the input.

In the absence of sensory stimuli, *I*_stim_ = 0, the ring network can show different spatial activity profiles depending on the form of the connectivity profile and the global excitatory drive *I*_exc_. For *I*_exc_ ≤ 0 the network has a unique equilibrium solution with zero activity, *r* (*θ*) = 0 Hz. Increasing the excitatory drive, *I*_exc_ > 0, will increase the activity of this equilibrium solution uniformly along the ring *r* (*θ*) = *r*_∞_ = Φ (*τw*_0_*r*_∞_ + *I*_exc_) /*τ* > 0. By studying the linear stability of this solution we can find a critical value of the global excitatory drive, *I*_exc_ = *I*_crit_, for which the spatially homogeneous state destabilizes into a spatially inhomogeneous state via a pattern forming instability known as Turing bifurcation. This new stable state will have a single bump of activity as shown in Fig. 1b for a connectivity profile that satisfies *w*_1_ > *w_k_* for all *k* ≠ 1. For our choice of the transfer function (Eq. 5) and the connectivity profile (see Tab. 1) this critical value is given by

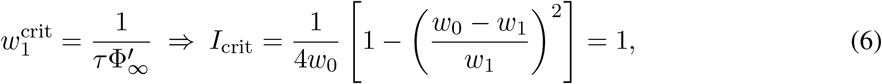

where 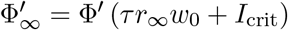 is the slope of the transfer function evaluated at the bifurcation point.

### Amplitude equation

The bump solution described in the previous section is analytically intractable. However, we can study its dynamics near the onset of the bump, i.e. in the vicinity of the bifurcation point *I*_exc_ = *I*_crit_, by performing a multiple-scale analysis based on a perturbation method^63^. We chose *I*_exc_ as our bifurcation parameter and define the perturbation

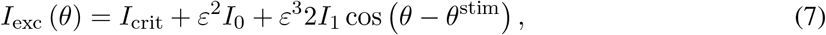

and the corresponding expansion of the firing rate

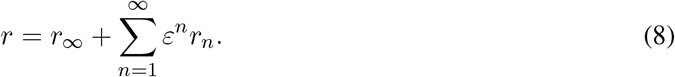

The multiple scale method assumes that the dynamics of the network can be separated into two independent time scales, such that we can define the slow time *T* = *ε*^2^*t* in which the dynamics of the bump evolve. Plugging all these definitions into the equation of the ring network, Eq. 4, we obtain the equation that describes the evolution of the amplitude of the bump, *A*(*t*), close to the bifurcation point:

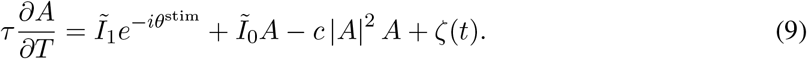

where 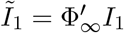 is the scaled stimulus strength, 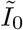 corresponds to the scaled distance to the bifurcation point, which is related to the global excitatory drive:

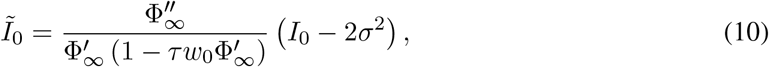

and the cubic factor *c* is

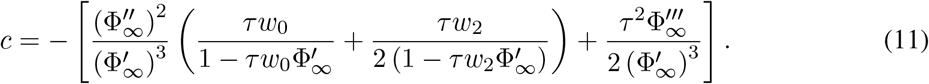

The last term in eq. 9 corresponds to the first Fourier component of the noise process *ξ*(*θ, t*) in Eq. 4, that can be written as

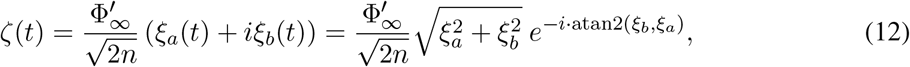

where, *n* is the discrete number of Fourier modes of the noise process *ξ*(*θ, t*), and *ξ_a_*(*t*) and *ξ_b_*(*t*) are independent Ornstein-Uhlenbeck processes with the same amplitude, *σ*_OU_, and time constant, *τ*_OU_ as *ξ*(*θ, t*). Additionally, the non-linear filtering of the noise process, *ξ*(*θ, t*), through the transfer function, introduces spatial correlations^64,65^ that effectively shift the bifurcation point to 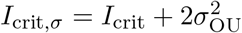, which explains the correction term in Eq. 10. Thus, changing the noise level not only increases the variability of responses (Fig. 4c), but also affects the integration of the stimulus by modulating the amplitude of the bump in a similar way as the global excitatory drive, *I*_0_.

Finally, by considering the polar form of *A*(*t*) = *Re*^-*iψ*^, Eq. 9 results in the system described by Eqs. 1a and 1b with the noise terms *ξ*_1_ and *ξ*_2_ being

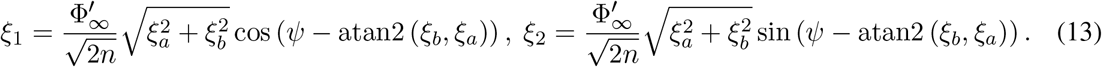

The values of the parameters used for the simulation of the amplitude equation are listed in Tab. 2.

**Table 2.** List of the parameters of the amplitude equation, and parameters of the numerical integration, used to simulate both Eqs. 1 and 4.

| Amplitude equation (Eq. 1) | Symbol | Value |
| --- | --- | --- |
| Additional excitatory drive, $I_{\text{exc}} - I_{\text{crit}}$ | $I_0$ | $\in [0.01, 0.2]$ |
| Slope of $\Phi(\cdot)$ at $I_{\text{exc}} = I_{\text{crit}}$ | $\Phi'_{\infty}$ | 50 |
| 2nd derivative of $\Phi(\cdot)$ at $I_{\text{exc}} = I_{\text{crit}}$ | $\Phi''_{\infty}$ | 100 |
| 3rd derivative of $\Phi(\cdot)$ at $I_{\text{exc}} = I_{\text{crit}}$ | $\Phi'''_{\infty}$ | 0 |
| 1st order prefactor | $\tilde{I}_0$ | $\in [-7/300, 31/300]$ |
| Cubic prefactor | $c$ | 1/3750 |
| Scaled stimulus amplitude | $\tilde{I}_1$ | 0.125 |
| Noise terms | $\xi_i(t)$,<br>$i=1,2$ | Eq. 13 |
| Numerical integration | Symbol | Value |
| Integration time step | $\Delta t$ | 0.2 ms |
| Spatial discretization | $n$ | 200 |
| Number of trials | $K$ | $10^5$ |

The amplitude equation (Eq. 9) derived here corresponds to the normal form of a Turing bifurcation where the first spatial mode of the homogeneous state becomes unstable. Similar derivations can be found in the literature (see for example, refs.^57,58^ and ref.^27^ section 8.1.2), which also study the general case of separated excitatory-inhibitory populations. This low-dimensional description of the network accurately describes the bump dynamics close to the bifurcation point where the bump emerges, and it is qualitatively valid for a wide parameter regime (Fig. 3b), allowing us to study the dynamics of the bump analytically.

#### Dynamics of the phase for a stationary bump amplitude

An analytical expression for the approximate evolution of the phase, *ψ*(*t*), can be obtain by assuming an almost static bump amplitude 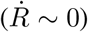, and then solving the ordinary differential equation Eq. 1b,

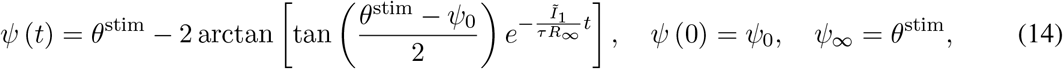

where *ψ*_0_ is the initial phase of the bump, and the stationary amplitude, *R*_∞_, is obtained by solving the third order polynomial that results from equating Eq. 1a to zero. The solution to this polynomial equation depends on the value of 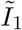, which if set to 0, gives 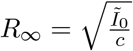. Extended Data Fig. 2a shows the evolution of the phase of the bump for an initial condition in which the bump is completely formed and centered at 0° for a stimulus *θ*^stim^ = 90°. The approximation (shaded lines), Eq. 14, is plotted along with the corresponding simulation of the amplitude equation showing good agreement. As expected, the approximation works better for larger values of 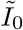, that make the amplitude of the bump less sensitive to the stimulus.

A first order approximation of Eq. 14, as a function of *θ*^stim^ (Extended Data Fig. 2c), is given by

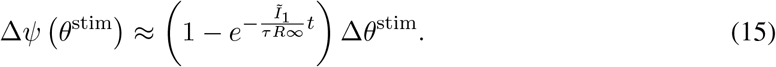

The error incurred by this approximation is small (approx. 10% for Δ*θ*^stim^ = 90°, *t* = 250 ms and 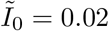), confirming that stimulus integration in the bump attractor model is relatively linear across a wide range of Δ*θ*^stim^.

#### Potential of the amplitude equation

Here we show that the system 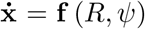 describing the evolution of the bump (Eq. 1), is indeed a gradient system, 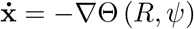, with a compatible potential function Θ (*R, ψ*) given by Eq. 2:

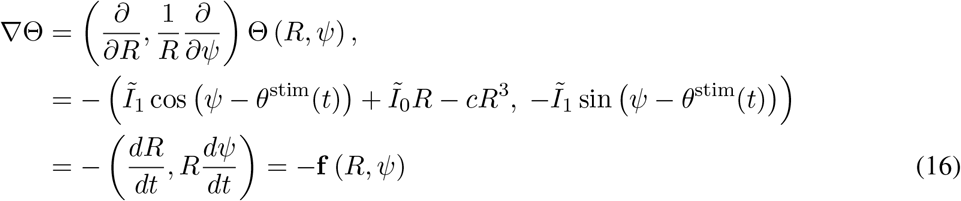

where we have used the polar form of the gradient operator and the differential of **x** = (*R, ψ*) is (*dR, Rdψ*). Similarly, we can compute its time-evolution

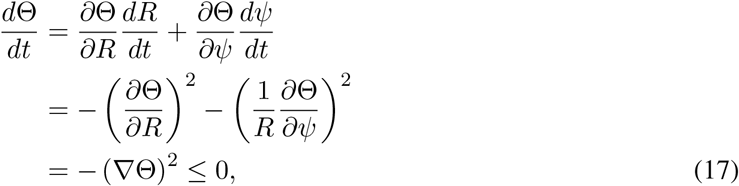

that shows that for all initial conditions **x**_0_ = (*R*_0_, *ψ*_0_), the potential will always decrease towards a stationary stable state (*R*_∞_, *ψ*_∞_).

### Dynamical system for the cumulative circular average (perfect vector integrator)

In order to understand the integration process carried out by the ring attractor model, we derived a dynamical system that perfectly integrates a circular variable to compute the circular average. In general, the optimal strategy to keep track of the average, 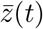, of a time-varying stimulus, *z*(*t*), is to compute the cumulative running average. For discrete-time stimuli the cumulative running average can be computed by the following iterative equation:

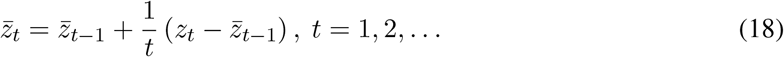

where *t* represents discrete time points. If we assume that the updates 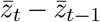 are small enough, we can write the above equation as a continuous process in time:

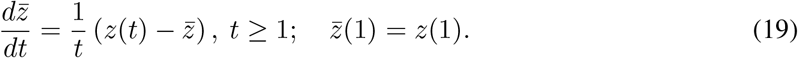

In order to avoid dealing with a non-autonomous system we can rewrite this equation by absorbing the time dependence into a new variable 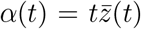 which leads to the integral form of the perfect integrator:

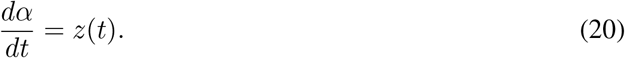

According to this equation, *α*(*t*) is just the sum of all *z*(*t*) (integral over time), and the average is obtained by dividing the summation by *t*:

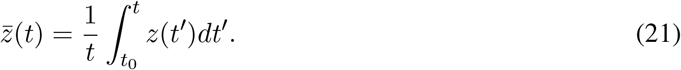

However, even if the result of the above integral is equivalent to that obtained by means of the iterative method, Eq. 19, the computational approach is fundamentally different. Here we would be required to memorize the entire time-varying stimulus up to the instant *t*, and divide its summation by the duration of the stimulus. In contrast, using the iterative method, Eq. 19, only requires keeping track of the average quantity at the last time point and update that value by weighting successive evidence in a time-dependent manner.

Eq. 20 holds for any time-varying process *z*(*t*), in particular, here we are interested in 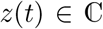. In this case we can interpret *z*(*t*) as stimulus vectors defined by the stimulus strength and the stimulus direction and 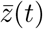 as the vector average up to time *t*. The resulting complex quantities can be written in polar form as *α*(*t*) = *R*(*t*)*e*^*iψ*(*t*)^ and *z*(*t*) = *I*(*t*)*e*^*iθ*(*t*)^ which gives the two equations

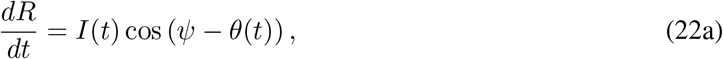

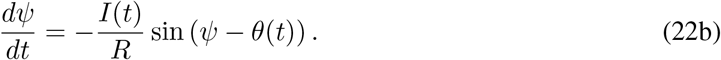

We refer to these equations as the *perfect vector integrator (PVI)* (Extended Data Fig. 3a,c). As described previously in this work, our main goal is to compute the circular average of the stimulus orientations, i.e. the argument of 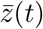. From the definition of 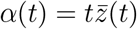, we can already see that 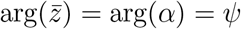. Therefore Eq. 22b exactly describes the evolution of the circular average of the orientations of the stimulus frames, i.e. the PVI computes the cumulative circular average. Moreover, note that Eq. 22b will compute the circular cumulative running average for any given normalization function *f*(*t*) such that 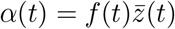.

The variable *R*, needed in the computation of the circular average, has several interpretations. First, the average magnitude of 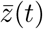, can be computed as 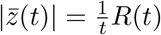. In other words, *R*(*t*) is the magnitude of the integrated vectors (i.e., the length of the resulting vector). Second, from this geometric interpretation it is easy to see that *R*(*t*) measures the dispersion of the directions *θ*, with *R*(*t*) growing linearly with *t* if *θ*(*t*) is a constant. If *θ*(*t*) is sampled from some underlying distribution, the growth will be slower and depends on the width of the distribution (Extended Data Fig. 3b,d). The circular variance of the samples can be computed from the ratio of *R*(*t*) and the maximum *R* obtained for *θ*(*t*) = const. as 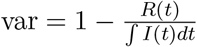. Third, if we assume that *θ*(*t*) is sampled from a von Mises distribution with mean *μ* and concentration *κ*, the running circular mean *ψ*(*t*) will also be distributed according to a von Mises distribution, with concentration proportional to *R*(*t*)*κ*. In a Bayesian framework, *ψ*(*t*) and *R*(*t*) thus track the mean and the concentration of the posterior distribution^66^.

#### Potential landscape for the perfect vector integrator

We follow the same approach used in the main text to graphically picture the dynamics of the perfect vector integrator (Eqs. 22a and 22b) by defining its potential landscape,

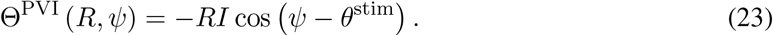

Proceeding in the same way as with the amplitude equation (Eq. 16) it is easy to see that the gradient of this potential gives the system Eq. 22.

### Categorical readout: decision circuit

In order to obtain a categorical decision from the continuous estimate of the ring network (Fig. 7), we use a standard two-population discrete attractor network^8,9,11^ that receives excitatory inputs from neurons located at each side of the ring (Fig. 7a). The decision circuit is composed of two recurrently coupled excitatory populations selective to left and right inputs, respectively, and an inhibitory population that is coupled to both excitatory populations. The activity of the network is described in terms of the firing rate of its populations, *r_L_, r_R_* and *r_I_*, and obeys^9^:

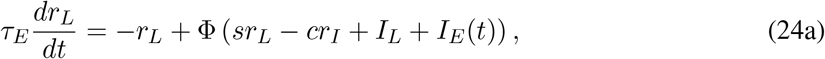

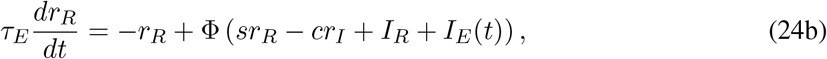

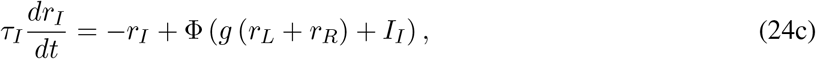

where *τ_E_*(*I*) represents the membrane time constant of the excitatory (inhibitory) neurons and 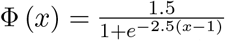 is the current-to-rate transfer function^9^. The inhibitory population receives excitatory inputs from both excitatory populations with synaptic strength *g*, and a constant positive current *I_I_*. The excitatory populations are recurrently coupled with synaptic strength *s* and receive inhibition from the inhibitory population with synaptic strength *c*. The values of all model parameters are summarized in Tab. 3. Additionally the L and R populations receive excitatory inputs from the integration circuit, *I_L_* and *I_R_*, that will depend on the size and location of the bump, and are computed as the convolution of the activity of the bump and a spatially constant kernel targeting its corresponding side of the ring:

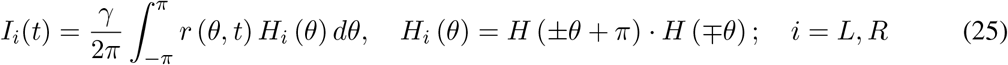

where *γ* = 0.0015, and *H_i_* (*θ*) is the left or right Heaviside step function.

Finally, *I_E_* represents a time varying external input that determines the dynamical regime of the decision network. During the integration period, *I_E_* = *I_R_* is set such that the activities of both populations remain at the low symmetrical state with bistability between the symmetric and the winner-take-all states. To force the decision, *I_E_* is increased by activating a transient urgency signal, *I_U_* (with duration Δ*T_U_*), that breaks the stability of the low-activity symmetric state and triggers the winner-take-all competition between the excitatory populations:

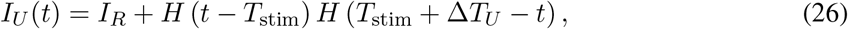

where *H*(*t*) is the Heaviside step function and *T*_stim_ is the stimulus duration. The decision is made whenever the activity of one of the excitatory populations crosses the 50 Hz threshold, *r_th_* (Fig. 7b,c).

**Table 3.** List of the parameters of the decision circuit and the urgency signal.

| **Decision circuit (Eq. 24)** | *Symbol* | Value |
| --- | --- | --- |
| Time constant of the excitatory/inhibitory neurons | $\tau_E, \tau_I$ | 20 ms |
| Strength of the recurrent coupling | $s$ | 1.9 |
| Strength of the inhibitory cross coupling | $c$ | 1.8 |
| Strength of the excitatory cross coupling | $g$ | 1 |
| External input to inhibitory neurons | $I_I$ | 0.2 |
| **External input, $I_E(t)$** | *Symbol* | Value |
| Resting input | $I_R$ | 0.33 |
| Resting input + Urgency signal | $I_R + I_U$ | 0.46 |
| Duration | $\Delta T_U$ | 500 ms |

### Modeling focused and divided attention

To model evidence integration in the focused vs. divided attention conditions (Extended Data Fig. 7) we built an extended network model consisting of two identical ring networks, each integrating one of the two spatially separated orientation streams (Extended Data Fig. 7d). The dynamics of both circuits are described by Eq. 4, using the parameters shown in Tab. 1. Focused and divided attention conditions are modeled by including a modulatory attention signal, in the form of an external input, that distributes a fixed amount of resources between the two circuits. For simplicity, the attention signal is included in the global excitatory drive, *I*_exc_ = *I*_crit_ + *I*_0_. In the focused condition 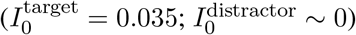 the resources are mainly directed towards the circuit encoding the target stream, while in the divided attention condition 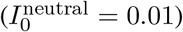 they are equally divided between both neutral circuits. Note that the amplitude of the bump goes as 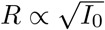, and therefore the overall activity in both conditions remains similar. The attention signal is the only difference between focused and divided attention trials. In particular, the stimulus strength is the same for cued and neutral stimulus streams. This is justified by the empirical finding that occipital and parietal EEG signals recorded from the participants represented neutral and target stimuli (in divided and focused attention, respectively) with a similar strength^41^. EEG signals also revealed that the information about the distractor stimulus is (partly) filtered out at early processing stages. Thus, in the model, the distractor stream in the focused attention condition has a 50% reduced stimulus strength and it is unattended 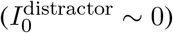. As a result, the distractor stream is not integrated, as has been observed in humans.

### Model simulations

#### Stimuli

The stimulus in each trial consists of a sequence of *N* i.i.d. oriented frames 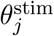, *j* = 1,…, *N* that are sampled from some probability distribution. We define the stimulus orientation as a piecewise constant function of time

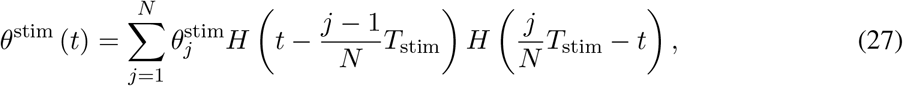

where *H*(*t*) is the Heaviside step function and *T*_stim_ is the stimulus duration. The input current to each neuron on the ring was modeled as

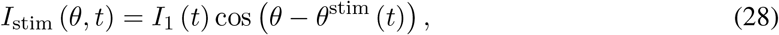

where *I*_1_ is the stimulus strength and *θ* is the position of the neuron on the ring (Eq. 4). In most of the simulations we run *K* = 10^5^ trials each consisting of *N* = 8 stimulus frames with a constant strength *I*_1_ and stimulus orientations sampled from a uniform distribution in the range 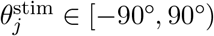, but randomly selecting combinations 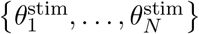 such that their circular mean, 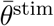, is uniformly distributed in the range 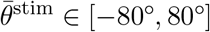.

#### Initial conditions

To simulate the **No initial bump** condition (Figs. 1c-e, 2b, 3b-f, 4 and 5a) the activity of the network at the beginning of each trial is set at a spatially homogeneous state (*I*_exc_ = *I*_crit_) with an average activity around *r*_∞_ = 12.5 Hz. The bump state is formed by increasing the global excitatory drive to the network, *I*_exc_, such that the bump starts to emerge at the beginning of the stimulus presentation.

On the other hand, for the **Initial bump** condition (Figs. 3a,c, 5b and 4b) the activity of the network at the beginning of each trial is set at a stationary bump. We compute this state taking into account both the strength of the stimulus, *I*_1_, and the effect of the noise (see the derivation of the Amplitude equation), by waiting long enough such that the system reaches the steady state. For the *biased* condition (Figs. 3a,c, and 5b), the bump is centered at 0°, while for the *unbiased* condition (Extended Data Fig. 4b), the initial phase of the bump is shifted to match the orientation of the first stimulus frame. This shift is taken to be equal to the position of the bump in the *No initial bump* condition at the end of the first stimulus frame, which still captures the impact of fluctuations at the beginning of the trial.

#### Numerical integration

We simulated Eqs. 4 and 9 following the Euler-Maruyama scheme with a time step Δ*t* = 0.5 ms, and discretizing the angular space *θ* into *n* = 200 evenly distributed locations, *θ_k_* = *k* · 360°/*n* – 180°. All simulations and analysis were done in Python 3.7.3, using standard libraries including among others: `NumPy, ScyPy` and `Pandas`.

### Statistical measures and fits

#### Estimation bias and estimation error

To characterize the accuracy with which an estimate 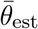 describes the true stimulus average 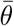, we define the following quantities.

##### Bias of the estimator

Usually, the estimation bias is defined as 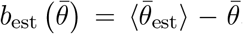, i.e., the difference of the trial-average of the estimates 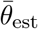 and the true value 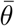. For our simulations (Fig. 5c), the bias defined in this way would be zero for 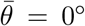 and increase approximately linearly with 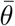. Thus, we preferred to define the estimation bias as one minus the slope of the estimation curve obtained from a regression fit (*k*_1_ in Eq. 32). An unbiased estimator would have a slope of 1 and thus zero bias (1 – *k*_1_ = 0), while slopes smaller than 1 correspond to increasingly large values of the estimation bias.

##### Variance of the estimator

The variance of the estimator is defined as 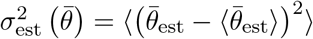, i.e. the variation of the estimates about their mean value. In Fig. 5e, we plot the square root of this quantity, the standard deviation of the estimates.

##### Estimation error

To assess the total deviations of estimated average orientations 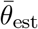 from the true stimulus average 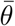, due to estimation bias and variance, we compute the root-mean-square error

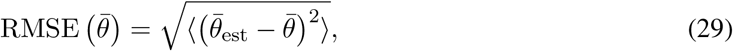

i.e., the square root of the trial-average squared estimation error. For Figs. 4c and 5e, we computed the mean of the RMSE across all average orientations 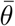.

##### Weighted circular average

To check whether other factors, besides the intrinsic nonlinearities of the model, are contributing to the estimation error, we compute the weighted circular average by simulating the PVI (Eq. 22), with the same exact stimulus *I*^stim^(*t*) used to simulate the ring network (Eqs. 27 and 28) but with a time-varying amplitude obeying

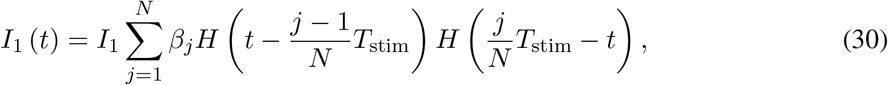

where *β_j_* correspond to the weights of the PK obtained for the ring network, and *H*(*t*) and *T*_stim_ are the Heaviside step function and the stimulus duration, respectively. This measure corresponds to the purple line in Fig. 4c.

#### Regression models for circular data

We use regression models to assess the impact of each stimulus frame on the final estimate of the stimulus average provided by the bump attractor model. Standard linear regression is not well suited for this purpose because we are estimating the average of a circular quantity. We thus derived a regression model for circular data. We define the weighted circular mean as the argument of a weighted sum of complex numbers:

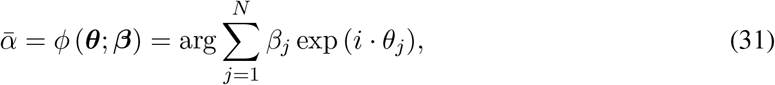

where *N* in the number of stimulus frames, *θ_j_* the orientation of the *j*th frame and *β_j_* the corresponding weight. In order to account for systematic deviations of estimated average orientations, we include a bias term *k*_0_ and a scale factor *k*_1_ (determining the slope of the estimation curve) and model the average stimulus estimation in trial *i* as:

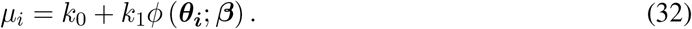

Assuming that the data follows a von Mises distribution with concentration parameter *κ*, we can derive the log-likelihood of observing the data *ψ* (i.e., the reported average angles) given the weights *β* as

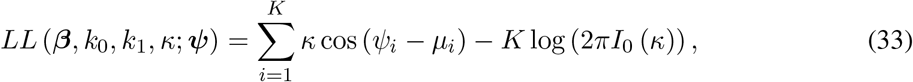

where *K* is the number of trials and *I*_0_ (*κ*) is the modified Bessel function of order 0. To uniquely define the weights *β* we include the constraint that the sum of ∑*β_j_* = 1. The parameters (*β, k*_0_, *k*_1_, *κ*) that maximize the log-likelihood were estimated using the `minimize` function from `scipy.optimize` module in Python 3.7.3.

#### Psychophysical kernel

For the model simulations, we obtain the PK as the stimulus weights of a circular regression model. To quantify the shape of a PK, we define the PK slope^54^. It is the slope of a linear regression of the PK, normalized between −1 (decaying PK, primacy) and +1 (increasing PK, recency). Formally, we first fit the PK with a linear function of time,

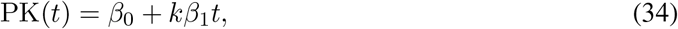

where *β*_1_ is the PK slope and 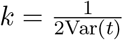 is a factor that normalizes the PK slope to the interval (−1,1).

### Psychophysical data and data analysis

#### Psychophysical experiment and datasets

We used published data from three studies carrying out psychophysical experiments in which human subjects had to categorize the average direction of a stream of Gabor patterns^6,7,41^ (Fig. 6a). To deal with slight differences in the experimental procedures (see below), we organized the data in 5 datasets: dataset 1 containing data from 15 subjects of Wyart et al.^6^, dataset 2 with data from 10 subjects of experiment 1 from Cheadle et al.^7^, dataset 3 with data from 19 subjects of experiment 2 from Cheadle et al.^7^, dataset 4 with data from 17 subjects of the focused attention condition from Wyart et al.^41^, and dataset 5 with data from the same 17 subjects in the divided attention condition Wyart et al. ^41^. The stimuli in all experiments consisted of a stream of eight oriented high-contrast (50%) Gabor patches with orientations 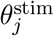, *j* = 1, 2,…, 8 uniformly distributed in the range between −90° and 90°. The duration of each sample was 250 ms in datasets 1-3 and 333 ms in dataset 4-5. In each trial participants reported whether, on average, the orientation of the eight samples fell closer to the cardinal or diagonal axes. The category level of each trial was computed by means of a folded decision-mapping rule consisting of a w-shaped profile that mapped the orientation of each sample to a decision variable ranging from −1 (diagonal) to 1 (cardinal), and then averaged across samples. Participants responded by pressing, with their left or right index finger, either two keys of a standard keyboard (dataset 1) or a one of two response buttons (DirectIN high-speed button box, Empirisoft; datasets 3-5). In the experiments of dataset 2 they responded at a continuous scale by clicking the mouse to a position on the screen corresponding to the integrated decision value. Auditory feedback was given at the end of each trial depending on the agreement between the response and the sign of the average decision value. The average number of trials performed by each participant was 546, 387, 433, 172 and 182 in datasets 1-5, respectively.

#### Analysis of psychophysical kernels for behavioral data

In this analysis (Fig. 6b) we included data from 61 subjects (datasets 1-4). Dataset 5 was not included because of the different experimental condition (divided attention that leads to more recency^41^). PKs of each participant are obtained as the weights of logistic regression (Extended Data Fig. 6). To characterize the shape of individual PKs, we used logistic regression with the constraint that the PK is either uniform (constant) or a linear function of time (Extended Data Fig. 6). We compare the model fits using AIC and classify each PK as uniform (best model has a constant PK), primacy (linear model with negative slope) or recency (linear model with positive slope).

#### Analysis of reaction times

In this analysis (Fig. 7) we included data from 47 subjects from datasets 1, 3 and 4. Dataset 2 was not included because subjects had to move and click the mouse which is not comparable to pressing a key or a button as in the other experiments. In addition, 4 subjects from dataset 3 were excluded because they had reaction times shorter than 240 ms in some trials and those reaction times had not been recorded. To compute the dependence of reaction times on the PK slope (Fig. 7f), we normalized the mean reaction times of each subject across the different datasets by z-scoring each dataset individually. This removes the overall differences in reaction times between experiments with a standard keyboard (dataset 1) and response buttons (dataset 3 and 4).

#### Analysis of evidence integration under focused and divided attention

For this analysis (Extended Data Fig. 7), we included data from the 17 subjects of dataset 4 and 5.

## Code availability

The code to generate the figures of the paper will be uploaded in github.

## Acknowledgements

We thank Valentin Wyarth and Christopher Summerfield for sharing the experimental data and for fruitful discussions, Tobias H. Donner, Bharath Chandra Talluri and Federico Devalle for excellent discussions, and Albert Compte, Joao Barbosa and Genis Prat-Ortega for comments on the manuscript. The research leading to these results has received funding from the Spanish Ministry of Science and Innovation together with the European Regional Development Fund (RYC-2015-17236 and BFU2017-86026-R to K.W, MTM2015-71509-C2-1-R and RTI2018-097570-B-I00 to A.R.) and from the Generalitat de Catalunya (grant AGAUR 2017 SGR 1565 to A.R. and K.W.).

---

**Extended Data Fig. 1.**
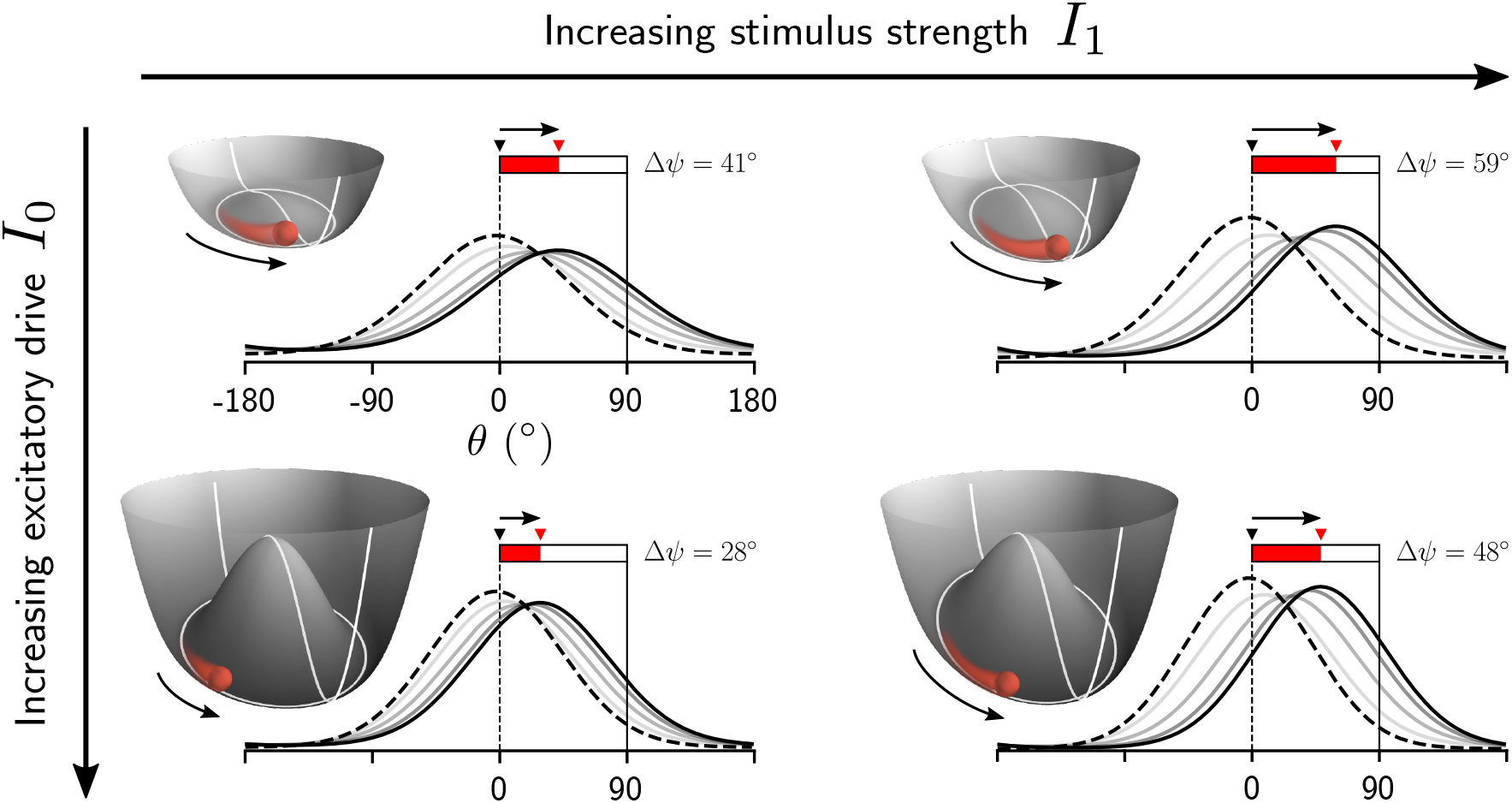
The global excitatory drive to the network and the stimulus strength determine the speed of movement of the activity bump. Displacement of a bump centered at 0° in response to a stimulus with average direction of 90° and duration of 250 ms (partially filled bar above the activity profiles), for different values of the excitatory drive and input strength. Mathematically, the angular speed of the bump is described by 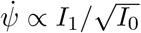. This relationship can be understood by representing the network activity as the movement of a particle in the potential (Eq. 2) corresponding to the amplitude equation, Eq. 1. On the one hand, increasing the excitatory drive to the network, *I*_0_, deepens the potential and increases the radius of the stable manifold (white circle), which in turn elongates the path that the particle has to travel in order to achieve a given angular displacement. Therefore, for the same stimulus strength, increasing the excitatory drive results in smaller displacements of the bump. On the other hand, increasing the stimulus input I_1_ has the opposite effect, i.e., the stimulus leads to a larger angular displacement for a given *I*_0_.

**Extended Data Fig. 2.**
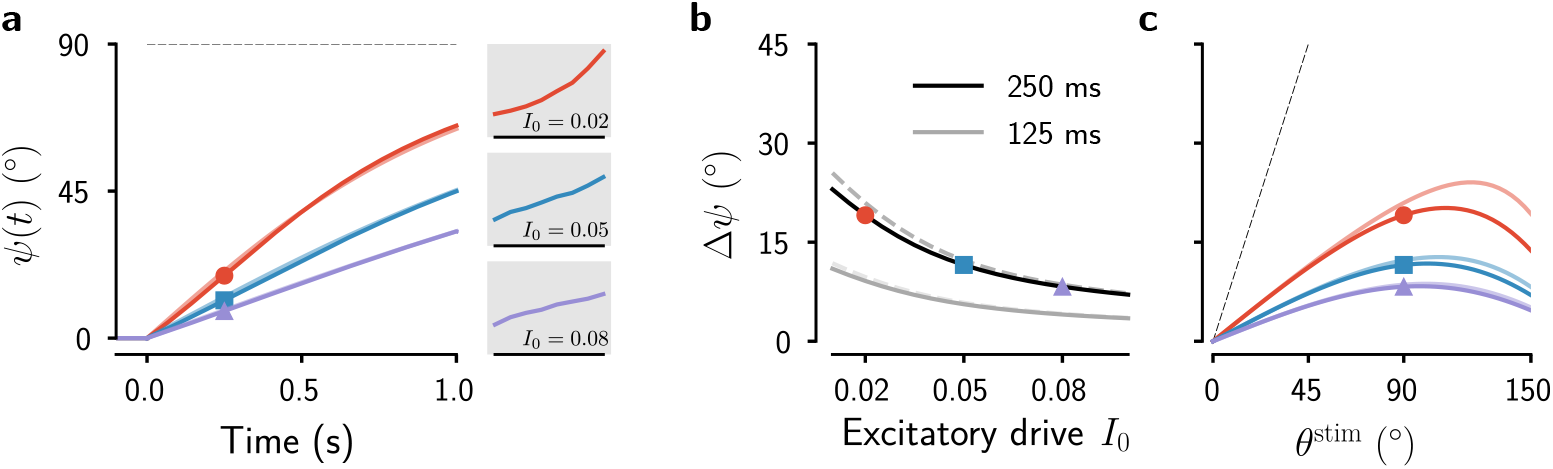
Impact of a single stimulus frame on the translation of the fully formed bump. **a**, Evolution of the phase of the bump for a single stimulus frame of strength *I*_1_ = 0.005 and duration *T*_stim_ = 1 s centered at *θ*^stim^ = 90°, for three different values of the excitatory drive *I*_0_. The solid lines correspond to simulations of the amplitude equation, Eq. 1, while the superimposed translucent lines have been analytically obtained using Eq. 14 which assumes a stationary bump amplitude. At the beginning of the trial the bump is fully formed and centered at 0°. Insets: PKs for each value of the excitatory drive (same as Fig. 3a). **b**, Change of phase as a function of the global excitatory drive to the network for a stimulus as in (a) but with a duration of *T*_stim_ = 250 ms (solid black) and *T*_stim_ = 125 ms (solid gray). The dashed lines correspond to the analytical approximation Eq. 14. **c**, Change of the phase of the bump as a function of the stimulus direction for *T*_stim_ = 250 ms. The dotted line indicates a bump displacement equal to the stimulus direction.

**Extended Data Fig. 3.**
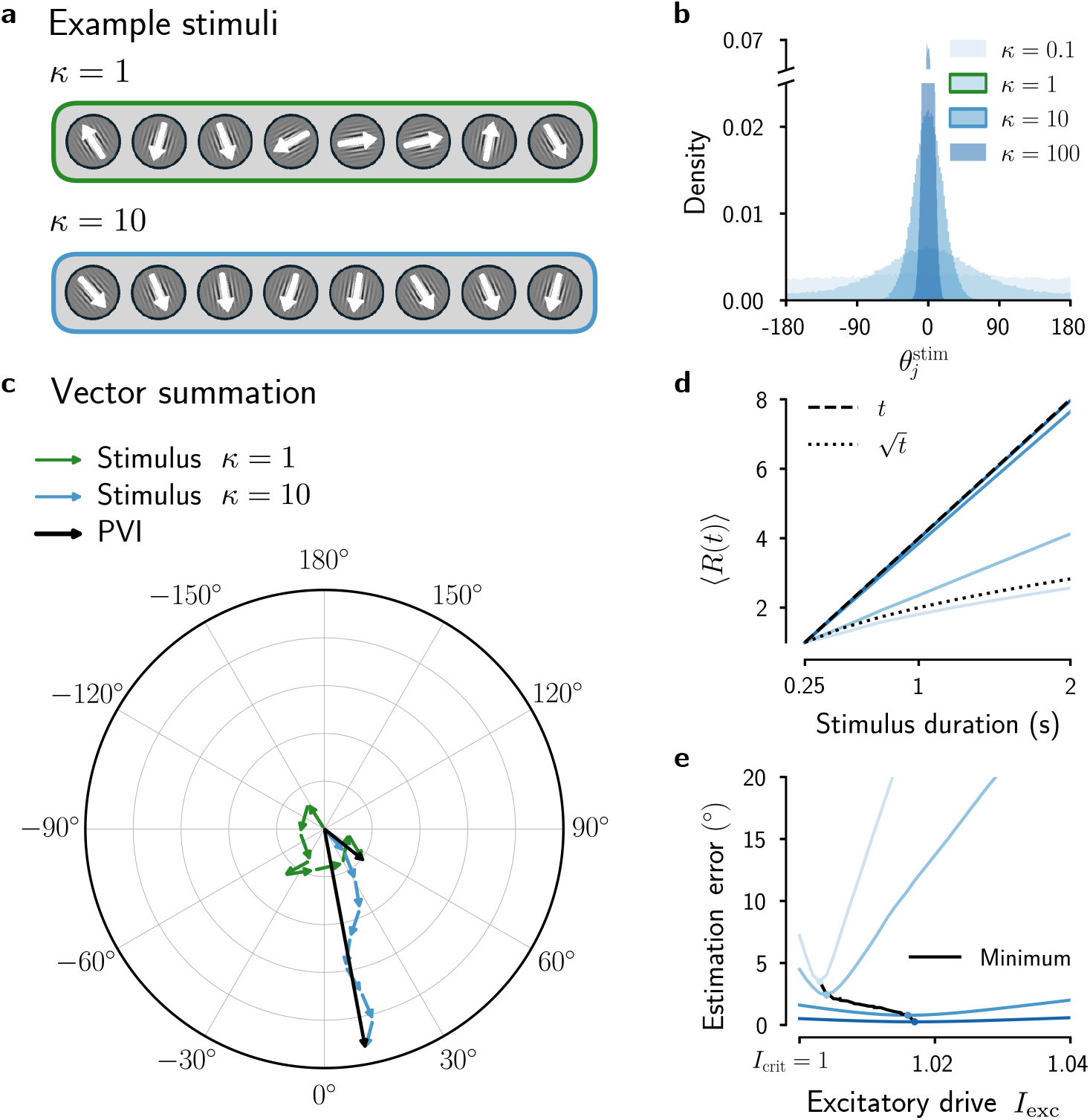
Perfect Vector Integrator (PVI) and the effect of the stimulus distribution on the estimation error of the bump attractor network. **a**-**b**, Example stimuli (**a**) with directions of individual frames, 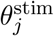 (indicated by white arrows), sampled from von Mises distributions with zero mean and different concentration parameter *κ* (**b**). **c**, Illustration of stimulus integration with vector summation for the two example stimuli from **a**. Each stimulus frame corresponds to a vector of length *I*_1_ (stimulus strength) and direction 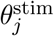 (green and blue arrows; see Methods, Eqs. 27 and 28). The direction of the vector sum (black arrows) gives the circular average of the stimulus directions 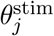. This is exactly the operation carried out by the PVI (Eq. 22), with the length of the vector corresponding to the modulus *R*, and the direction of the vector corresponding to the phase *ψ*. **d**, Evolution of the trial-averaged modulus 〈*R*(*t*)〉 of the PVI (Eq. 22a) for increasing stimulus duration. Stimuli had a fixed strength (*I*_1_ = 1), consisted of frames with 250 ms duration, and directions 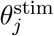 sampled from the distributions shown in **b**. When the concentration parameter *κ* is large, i.e., the dispersion of the stimulus directions is small, 〈*R*(*t*)〉 grows approximately linearly in time (as in the example shown in **c**, blue arrows). On the other hand, when the dispersion is large (e.g. *κ* = 0.1), 〈*R*(*t*)〉 grows much slower (as in the example shown in **c**, green arrows). A growth with 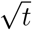 is shown for comparison. **e**, The root-mean-square estimation error (Methods) of the bump attractor model as a function of the global excitatory drive *I*_exc_ for the stimulus distributions from **b**. Simulations were performed using the amplitude equation (Eq. 1) with the noise term set to zero (*σ***ou** = 0). The optimal value of the excitatory drive *I*_exc_ to achieve the minimum error depends on the stimulus distribution (black line): For small values of *κ* (large dispersion), the model performs best close to the bifurcation point *I*_crit_. As *κ* increases (small dispersion), the optimal value of *I*_exc_ increases. A larger value of *I*_exc_ is beneficial in this case because it allows for larger bump amplitudes needed for the integration of stimulus frames from a narrow distribution (as in **d**). This holds even though larger *I*_exc_ increasingly lead to a juice-squeezer-shaped potential that deviates from the flat potential of the PVI (Fig. 3a). Note that *I*_0_ = *I*_exc_ – *I*_crit_ (Methods).

**Extended Data Fig. 4.**
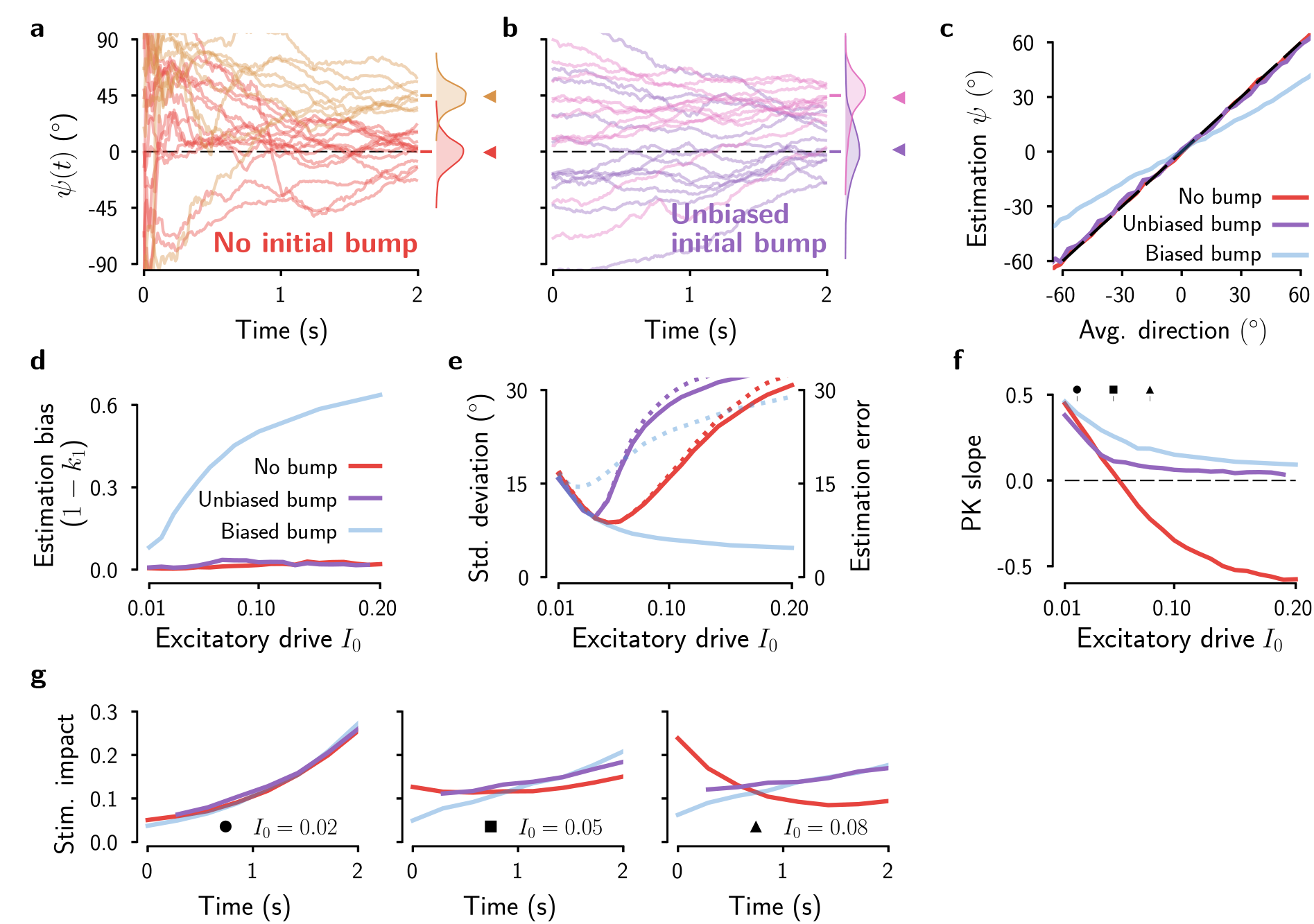
Bias and variance of the stimulus estimates with an initially formed bump and unbiased initial condition. **a** - **e**, as in Fig. 5 but instead of initializing the bump at 0° in the fully formed condition, its initial position is now determined by the direction of the first stimulus frame (**b**; Methods). **a**, Bump phase in example trials when starting with homogeneous network activity, re-plotted from Fig. 5a. **c**, Continuous estimations as a function of the mean stimulus direction. **d**, Systematic bias of the estimations as a function of the global excitatory drive *I*_0_. **e**, Standard deviation (solid lines, left axis) and root-mean-square estimation error (dotted lines, right axis) of the final bump position as a function of the excitatory drive *I*_0_. **f**, Slope of the psychophysical kernel versus the global excitatory drive *I*_0_. **g**, PKs (Methods) for the three conditions: no initial bump condition (red), initially biased bump condition (faint blue), and initially unbiased bump condition (purple). To compute the PKs of the initially unbiased condition the contribution of the first stimulus frame has been discarded. In sum, the figure confirms that the fixed initial bump condition, but not a fully formed bump per se, leads to an estimation bias. Compared to the *no initial bump* condition, the *unbiased initial bump* condition leads to a higher variance of the estimates and recency or uniform (but not primacy) PKs.

**Extended Data Fig. 5.**
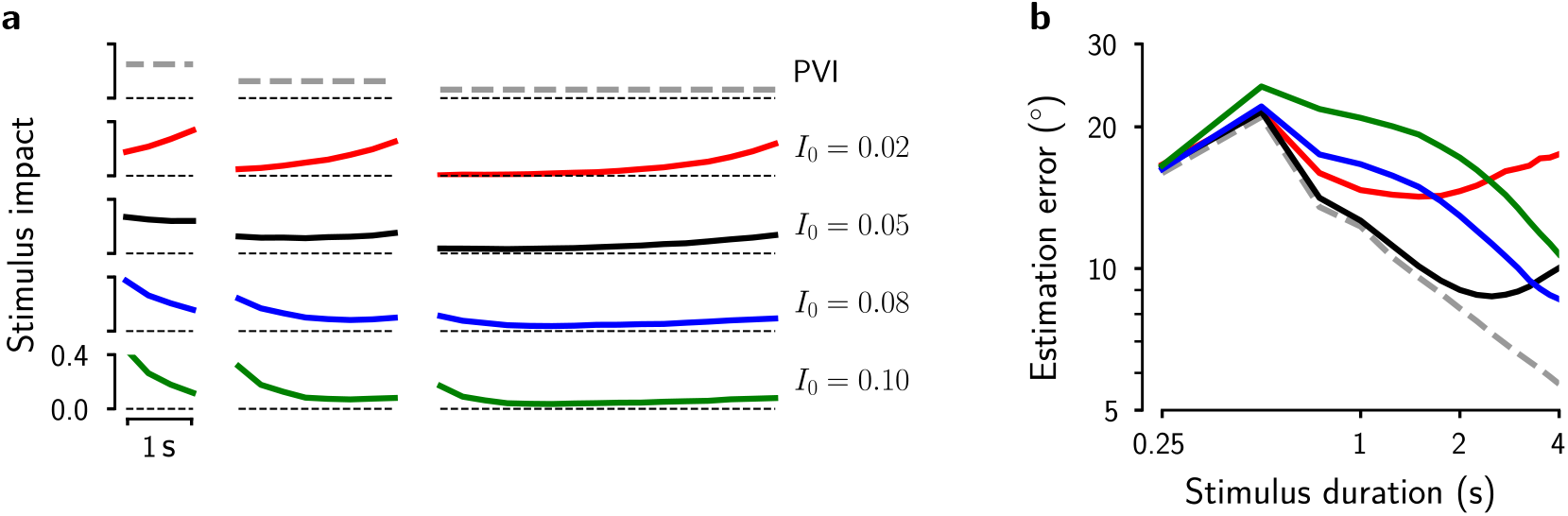
Dynamics of evidence integration in the bump attractor model for increasing stimulus duration. **a**, PKs of the PVI and the bump attractor model for increasing stimulus durations (*T*_stim_ = 1, 2 and 4 seconds) for different values of the excitatory drive *I*_0_. Stimuli were composed of frames with a fixed duration of 250 ms. The slopes corresponding to these PKs are shown in Fig. 3e. **b**, Root-mean-square estimation error (Methods) vs. stimulus duration for different levels of the excitatory drive (as in **a**). Frame duration was 250 ms. For comparison, we plot the estimation error of the PVI (Eq. 22; dashed line) to which we added noise terms obtained using Eq. 12. The estimation error in the PVI is entirely caused by this noise because the integration process itself is optimal. The particular form of the error evolution depends on the distribution of the stimulus orientations, e.g. for a pair of oppositely directed vectors the resulting vector is null and its direction is completely determined by the noise term. In general, the average modulus of the summation of two vectors will be widely distributed, and thus the fluctuation term determines the average orientation for a significant part of the distribution. This explains the initial increase of the estimation error, with a peak at 0.5 s (two stimulus frames). In contrast, the increased estimation errors—with respect the PVI—of the bump attractor model (colored lines) are due to the intrinsic dynamics of the network, and therefore depend on the integration regime.

**Extended Data Fig. 6.**
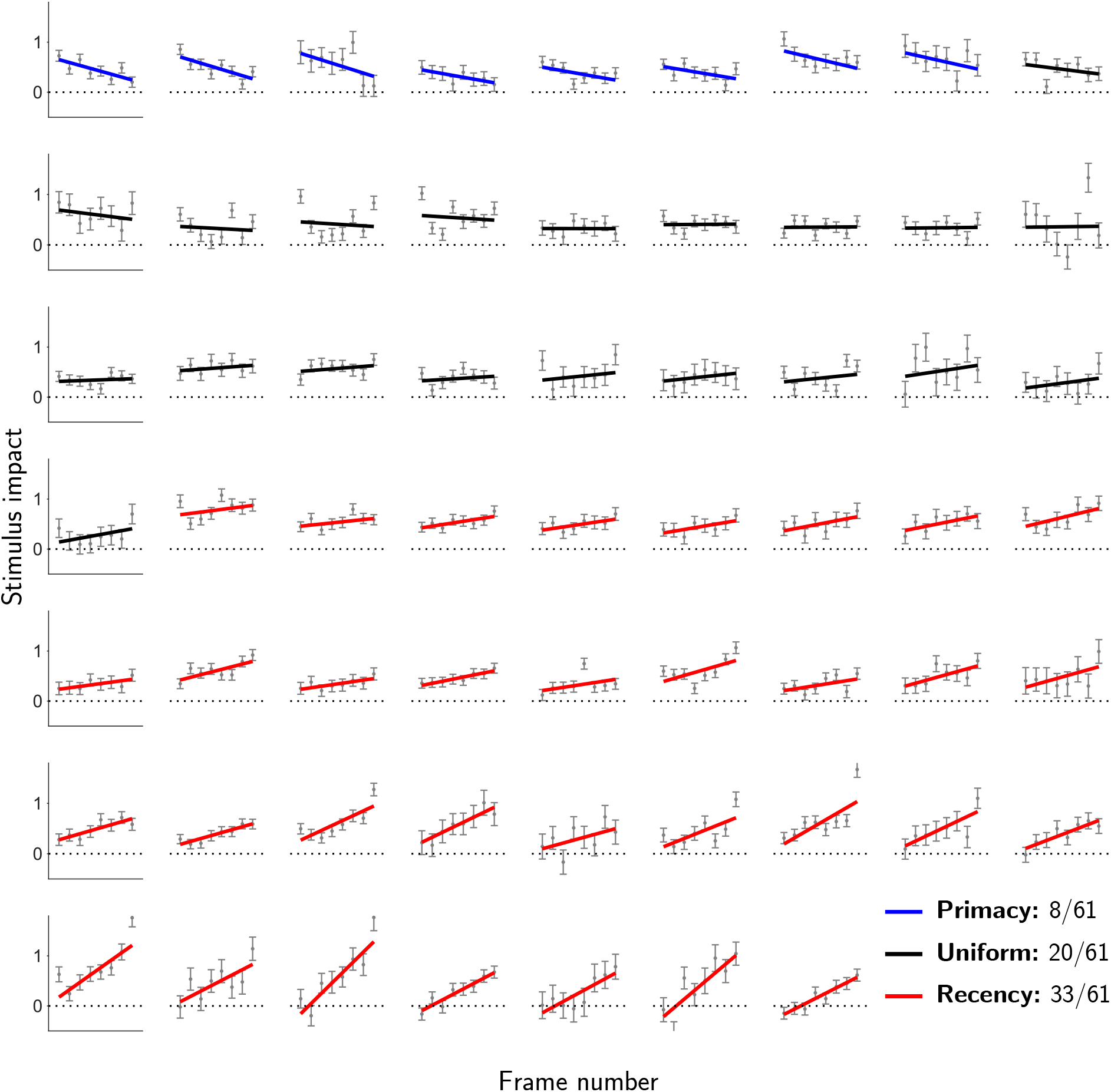
Individual psychophysical kernels from human subjects. Psychophysical kernels of individual subjects^6,7,41^. Colored lines are regression weights obtained from logistic regression with the constraint that the PK is a linear function of time (these fits are used to compute the PK slope). Plots are sorted and color coded by kernel shape. Gray errorbars show the weights obtained from logistic regression fits without applying constraints.

**Extended Data Fig. 7.**
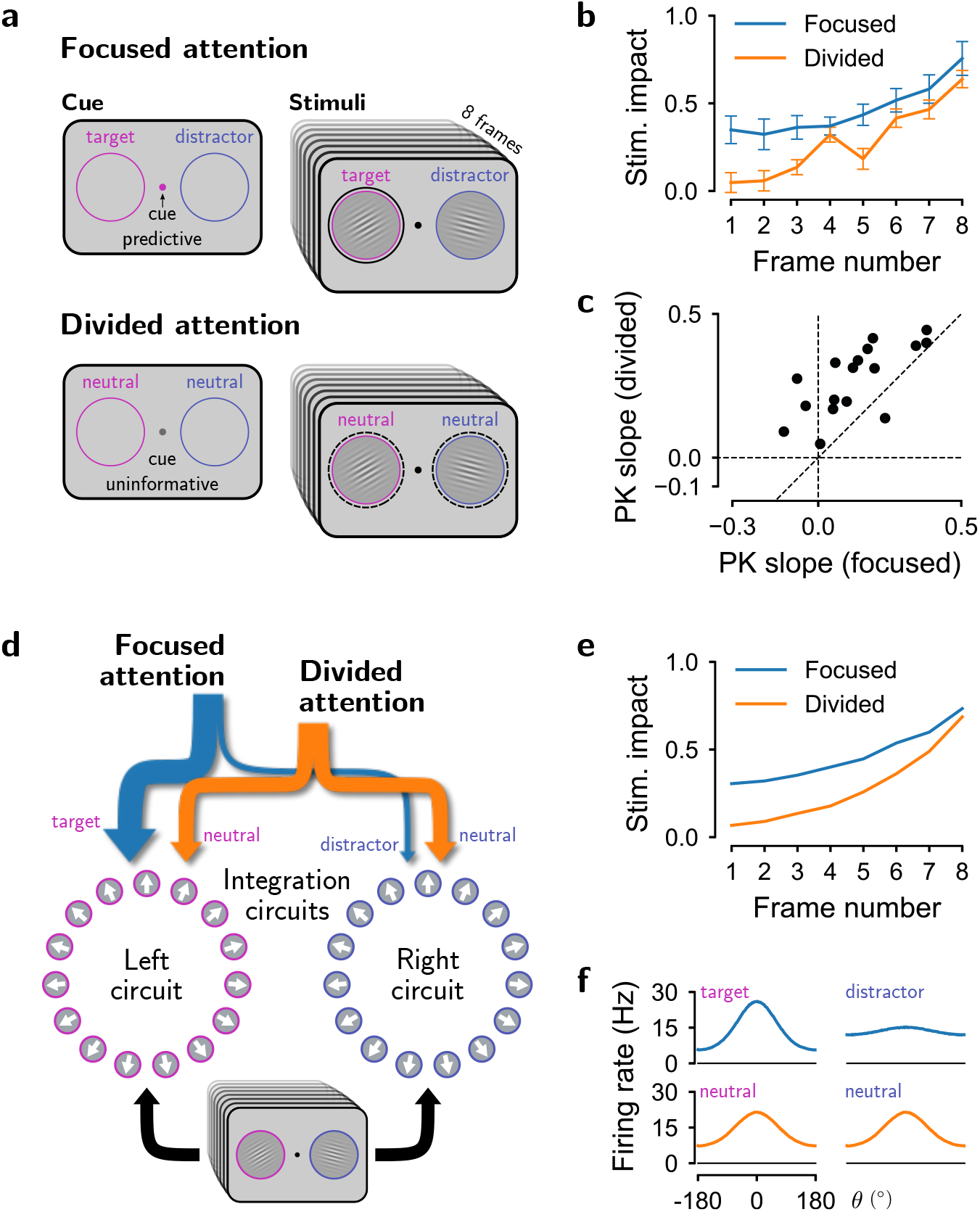
Evidence integration under focused and divided attention. As a further test of the explicative power of the ring attractor model, we investigated whether it could shed light on the neural mechanisms underlying a change in temporal evidence weighting when attention is divided between spatially separated information sources. **a**, Category-level averaging task with focused and divided attention conditions^41^. The task is similar as in Fig. 6a, but with two streams of visual gratings presented simultaneously. In the focused attention condition, the relevant stream is indicated by a cue in the beginning of the trial whereas in the divided attention condition both streams need to be integrated and one of them will be probed randomly. **b**, PKs for the relevant stream in the two attention conditions (data re-analyzed from^41^). Errorbars indicate the SEM across *n* = 17 participants. **c**, PK slope in the divided attention condition is consistently higher than in the focused attention condition (except for 1 / 17 subjects). This shows that humans can integrate information from two streams of visual gratings simultaneously, though with an increased integration leak, that is, with PKs that show more recency effect compared to the focused attention condition in which only one stream needs to be integrated. **d**, Model architecture, with two ring circuits that integrate two spatially distinct sensory streams independently in parallel. The strength of the stimulus input is the same across focused and divided attention conditions (Methods). The integration dynamics of the circuits is controlled by sustained attention modulation signals (blue and orange arrows, respectively) that effectively add to the global excitatory drive (*I*_0_) of the corresponding circuit. **e**, Model PKs for focused and divided attention conditions. We parameterized the model such that in the absence of the attention signal it operates in the recency regime. In the focused attention condition, the circuit integrating the target stimulus receives a large attention signal, increasing the total excitatory drive and pushing the network towards more uniform evidence integration (cf. Fig. 3c). In the divided attention condition, both circuits receive an attention signal of equal strength, albeit smaller than under focused attention. Therefore, inputs are now integrated with a more recency PK. **f**, Average population firing rate at the end of the trial for the target and the distractor stream in the focused attention condition (top) and for the neutral streams in the divided attention condition (bottom). This illustrates that the strong attention signal towards the circuit integrating the target stream in the focused attention condition led to a representation of the average stimulus orientation in a larger activity bump compared to the bumps produced in the divided attention condition. This is consistent with the experimental finding that the subject’s choice can be better decoded from preparatory EEG signals that reflect choice build-up in the focused attention condition (Fig. 6 in ref.^41^).

**Supplementary Video 1 I Single-trial activity of the ring network operating close to the perfect integration regime.** The **top left** panel shows the instantaneous activity of the network, with the red triangle marking the phase (center) of the bump that encodes the estimated average orientation of the stimuli. The true running average, computed by means of the perfect vector integration (PVI), is indicated by the blue triangle. At the beginning of the trial the amplitude of the bump is almost zero, and it grows and moves as the stimulus is presented. The **bottom left** panel shows the time-course of the position (phase) of the bump (red) and the PVI (blue) together with the stimulus stream on top. The light shaded region corresponds to the stimulus interval and the darker region marks the current stimulus frame. The black arrow marks the orientation of the current stimulus frame. As the simulation progresses the graph is updated to show eight additional stimulus frames. The **right panel** shows the potential (Eq. 2) corresponding to the network operating at approximately the uniform integration regime, with a global excitatory drive *I*_exc_ = 1.0055. The polar graph at the bottom corresponds to the axial projection of the potential with the red trace showing the state of the network in terms of the phase and the radius of the bump as described by the amplitude equation Eq. 1. The temporal evolution of the PVI is shown for comparison (blue trace). During the first two seconds (corresponding to eight stimulus frames) the network perfectly integrates the stimulus which is reflected on the equivalence between the phase of the bump and the perfect vector integrator. Once the bump amplitude cannot grow further due to the shape of the potential, the network starts to overweight new stimuli and thus shows a recency behavior.

**Supplementary Video 2 | Recency in the ring network.** As in Supplementary Video 1 but with a lower global excitatory drive *I*_exc_ = 1.002, that sets the network at the recency regime (see Fig. 3c,e). The maximal bump amplitude is small, limited by the form of the potential. As a consequence, the network overweighs the momentary stimulus information causing a constant leakage of the previously integrated sensory information, which results in a strong recency effect.

**Supplementary Video 3 | Primacy in the ring network.** As in Supplementary Video 1 but with a higher global excitatory drive *I*_exc_ = 1.01, that puts the network in the primacy regime (see Fig. 3c,e). During the first two seconds (eight orientation frames) the activity bump grows rapidly which causes an overweighing of early stimulus frames. However, once the amplitude of the bump saturates (Fig. 3d), the network will start to partly erase the accumulated information (leaky integration) exactly as in the recency regime (Supplementary Video 2).

